# Multiple Haplotype Reconstruction from Allele Frequency Data

**DOI:** 10.1101/2020.07.09.191924

**Authors:** Marta Pelizzola, Merle Behr, Housen Li, Axel Munk, Andreas Futschik

**Author notes:** These authors contributed equally (Marta Pelizzola, Merle Behr).

## Abstract

Since haplotype information is of widespread interest in biomedical applications, effort has been put into their reconstruction. Here, we propose a new, computationally efficient method, called `haploSep`, that is able to accurately infer major haplotypes and their frequencies just from multiple samples of allele frequency data. Our approach seems to be the first that is able to estimate more than one haplotype given such data. Even the accuracy of experimentally obtained allele frequencies can be improved by re-estimating them from our reconstructed haplotypes. From a methodological point of view, we model our problem as a multivariate regression problem where both the design matrix and the coefficient matrix are unknown. The design matrix, with 0/1 entries, models haplotypes and the columns of the coefficient matrix represent the frequencies of haplotypes, which are non-negative and sum up to one. We illustrate our method on simulated and real data focusing on experimental evolution and microbial data.

## 1 Introduction

Haplotypes are stretches of DNA that are inherited together from the same parent. They provide crucial information when analysing genetic data. Haplotype information is important to explore genetic factors affecting diseases [Tewhey et al., 2011], to impute missing genotype data [Marchini et al., 2007], to infer demographic population histories [Tishkoff et al., 1996], and to detect traces of selection [Sabeti et al., 2002]. Haplotypes can also be interpreted as microbial strains when studying for instance the gut microbiome of humans [Garud et al., 2019]. This is of considerable interest because of the association between the microbiome composition and several diseases, see, e.g., [Feng et al., 2015, Wang et al., 2012].

If sequence data are available on an individual basis at a sufficient read coverage, phasing methods are often used to infer haplotypes. Indeed, for human sequencing projects such as the 1000 genomes project [The 1000 Genomes Project Consortium, 2015], great efforts have been put into the development of fast algorithms to obtain haplotype information from diploid reads. In other fields of applications, however, this is not possible. In studies on the genetic basis of adaptation of non-human populations, for instance, the available haplotype information is often very limited, or lacking entirely. This is because either resources for sequencing are more scarce, or there are technological difficulties as for instance with micro-organisms. In order to lower the cost of the experiment, populations are frequently sequenced as a pool [Burke et al., 2010, Illingworth et al., 2012, Barghi et al., 2019]. This approach provides genome-wide allele frequency data at a SNP level [Futschik and Schlötterer, 2010, Schlötterer et al., 2014], but does not lead to any direct haplotype information.

Pool sequencing is frequently used for instance in Evolve and Resequence (E&R) experiments [Turner et al., 2011] that follow one or more (replicate) populations under controlled conditions and provide a time series of allele frequency data. Such experiments are carried out, since it is very challenging for natural populations to understand evolution at a genetic level because of the influence of unknown demographic factors and various selective pressures [Savolainen et al., 2013]. Even in the lab, ubiquitous polygenic adaptation leads to complex evolutionary patterns that can be better understood using haplotype information [Michalak et al., 2019, Karasov et al., 2010, Barghi et al., 2019, Burke, 2012]. Evolutionary phenomena are also investigated at a spatial level. [Meier et al., 2020] and [Jones et al., 2012], for instance, investigated *Heliconius* butterflies and sticklebacks, respectively, to understand adaptation from variation across hybrid zones. In these studies, the evolution of different species that originated from adaptive radiation ([Meier et al., 2020]) has been explored. In this context, haplotypes help to uncover the genetic basis of such adaptation processes. In medical applications the evolution of viruses within hosts is often of interest (see, e.g. [Zanini et al., 2015, Sudderuddin et al., 2020]). With longitudinal virus data from one (or more) given patients, each individual provides a pool of viral RNA which is sequenced through time. Haplotypes permit to understand the pattern of viral evolution both within and between hosts.

Given the large number of situations without direct haplotype information, efforts have been made to infer them from allele frequency data. With E&R in mind, the approach proposed in [Franssen et al., 2017] and optimized in [Otte and Schlötterer, 2019] reconstructs only a subset of SNPs that show a clear signal of selection (see Section 5 for further details). The methods by [Excoffier and Slatkin, 1995], [Pirinen, 2009], [Gasbarra et al., 2011], [Long et al., 2011], [Kessner et al., 2013] and [Cao and Sun, 2015] use known founder haplotypes, and only estimate the frequency trajectories of these haplotypes over time. To reconstruct microbial strains, methods have been proposed that rely on sequencing read data [Pulido-Tamayo et al., 2015], [Cao et al., 2020], [Knyazev et al., 2018].

In this paper, we propose a new principled approach, called haploSep, that is broadly applicable in situations where no candidate haplotypes from other sources are available. Indeed, it only requires allele frequency data from multiple samples as input. Using an iterative Lloyd’s type (see, e.g., [Lu and Zhou, 2016]) algorithm, it is computationally much faster than the above mentioned methods that rely on read data and can take days to run (see section S13 in the SI). Indeed, our procedure has a linear run time in the number of SNPs (potentially up to some initial clustering), and thus, can reconstruct several thousand SNPs for multiple haplotypes within a couple of seconds on a standard laptop.

In Section 2, we explain the proposed method in detail. In Section 3, we consider realistic scenarios that involve both real and simulated data to evaluate its performance. Conditions are also discussed under which we expect our approach to work. We also apply our method to four real data sets in Section 4. Our main focus is on applications to E&R experiments, but we additionally provide a real data example on HIV. For a comparison with a method that works on read data directly, see Section 5. We implemented our method *haploSep* in an *R* package available at https://github.com/MartaPelizzola/haploSep.

## 2 Methods

### Notation

In the following we use the notation [*N*] := {1*, …, N*} for an integer *N*. For a matrix *A* let *A*_*i*·_ and *A*_·*i*_ denote its *i*th row and column vector, respectively. With ǁ*A*ǁ and *A*^T^ we denote the Frobenius norm and the transpose of a matrix *A*, respectively. For a vector *a*, we always assume it is a column vector. We denote **1**= (1*, …,* 1)^T^ the vector with just ones.

Simultaneously reconstructing the structure of dominant haplotypes (a.k.a. major haplotypes) as well as their relative proportions in the population amounts to a matrix factorization problem with finite alphabet constraint on one of the matrices, and positivity as well as unit column-sums constraints on the other. More specifically, assume we obtained relative allele frequencies *Y* ∈ [0, 1]^*N*×*T*^ from a pool sequencing experiment, at time points *t* ∈ [*T*] and SNP locations *n* ∈ [*N*], from a population that consists of *m*_0_ haplotypes. Then the underlying population allele frequencies *F* ∈ [0, 1]^*N*×*T*^ can be written as

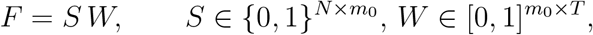

where *S*_·*i*_ ∈ {0, 1}^*N*^ for *i* ∈ [*m*_0_] denotes the genotype structure of haplotype *i*, that is, *S*_*ni*_ = 1 if haplotype *i* takes the reference allele at location *n* and *S*_*ni*_ = 0 otherwise. The frequencies *W*_*it*_ denote the relative proportion of haplotype *i* at time point *t* (haplotype frequency). Ignoring any sequencing error, we have *E*(*Y*|*F*) = *F.* Our aim is to reconstruct both the matrices *S* and *W* from the measurement matrix *Y*. This amounts to a specific type of a *finite alphabet blind separation* problem [Behr and Munk, 2017, Behr et al., 2018].

In general, *m*_0_, the overall number of haplotypes, can be very large, possibly *m*_0_ > *n*, which makes *S* and *W* non-identifiable, even from the noiseless allele frequencies *F*. However, when (for most of the time points *t*) the population is dominated by *m* ≪ *m*_0_ haplotypes, such that,

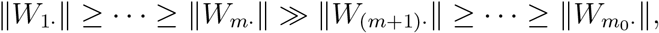

we denote structure and frequency of the dominant haplotypes as *S*^*d*^ = (*S*_*ni*_)_1≤*n*≤*N,*1≤*i*≤*m*_ and *W*^*d*^ = (*W*_*it*_)_1≤*i*≤*m,*1≤*t*≤*T*_ and obtain *F* = *S*^*d*^*W*^*d*^ + *B*, with a bias *B* = *SW S*^*d*^*W*^*d*^, which is the allele frequency component of minor haplotypes. In the following, we will omit the superscript *d* and just write

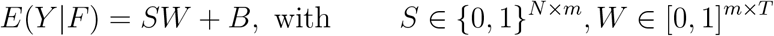

such that *m* ≪ *T, N* and ǁ*B*ǁ ≪ ǁ*SW*_·*i*_ǁ for *i* ∈ [*m*]. With our considered simulation and real data scenarios, a bias term only makes a difference when there is a large number of minor haplotypes present at several time points. For our further analysis we assume that the minor haplotypes *B* and the major haplotypes *SW* are independent and that on average, minor haplotype contributions *B*_*nt*_ only depends on the time points *t* ∈ [*T*], but not on the SNP location *n* ∈ [*N*]. This assumption is necessary in order to guarantee identifiability of the structure *S* and frequency *W* of the dominant haplotypes. Moreover, this assumption is justified for haplotype reconstruction as minor haplotypes will typically not contain selected positions and therefore are expected to show a similar average contribution among different SNPs. More precisely, we assume that

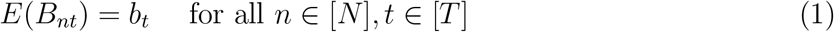

and thus,

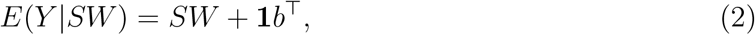

where *b* = (*b*_1_, …, *b*_*T*_)^T^ ∈ [0, 1]^*T*^ is the average bias term from the minor haplotype contribution.

### 2.1 HaploSep algorithm

If one had observed the allele frequencies *F* = *SW* + **1***b*^T^ without noise, it would be possible to uniquely recover *S, W* and *b* by exploring the ordering structure of the rows of *F* (assuming some w eak identifiability conditions on *S* and *W* as detailed in the SI). To this end, note that the *N* different row vectors of *S* (each of dimension *m*) can take at most 2^*m*^ different values, namely, the binary vectors in {0, 1}^*m*^. Thus, also the *N* different row vectors of *F* = *SW* + **1***b*^T^ (each of dimension *T*) can take at most 2^*m*^ different values. For example, when *m* = *T* = 2, those 2^*m*^ = 4 possible row vectors of *F* are (*b*_1_*, b*_2_), (*b*_1_ + *W*_11_*, b*_2_ + *W*_12_), (*b*_1_ + *W*_21_*, b*_2_ + *W*_22_), and (*b*_1_ +*W*_11_ +*W*_21_*, b*_2_ +*W*_12_ +*W*_22_). As *b*_*t*_, *W*_*it*_ ∈ [0, 1] and ǁ*W*_1·_ǁ ≥ ǁ*W*_2·_ǁ ≥ … ≥ ǁ*W*_*m·*_ǁ, it is easy to check that among those 2^*m*^ possible row vectors of *F*, the one with the smallest norm, *F*_*i*·_, corresponds to a haplotype structure where *S*_*i*_1·__ = (0, …, 0) and thus, *F*_*i*_1·__ = *b*, which allows to recover *b*. Similar, the second smallest row vector *F*_*i*_2·__ of *F* corresponds to the situation where *S*_*i*_2·__ = (0, …, 0, 1) and *F*_*i*_2·__ = *W*_*m*·_ + *b*, which allows to recover *W*_*m*·_. Proceeding in a recursive way, one can uniquely recover *S, W* and *b* from the noiseless matrix *F* = *SW* + **1***b*^T^. Details are given in the SI and pseudo code for this (noiseless) reconstruction algorithm is given in the `HaploSepCombiExact` (Algorithm 1 in the SI), see also [Behr and Munk, 2017].

However, in practice, one only obtains the noisy pool sequencing data *Y* but not the population allele frequencies *F*. Therefore, a direct application of `HaploSepCombiExact` is impossible and also not reasonable as further regulation will be required to obtain statistically stable estimates of *S, W* and *b*. Therefore, we consider a relaxation of the exact solution to *Y* = *SW* + *b*, i.e. we seek to solve the optimization problem

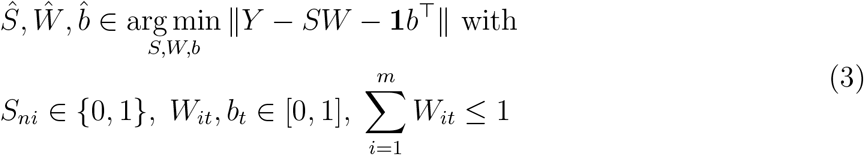

for *i* ∈ [*m*], *t* ∈ [*T*], *n* ∈ [*N*]. If *Y* were normally distributed, this would be the maximum likelihood estimator. For Pool-Seq data, due to its discrete structure, *Y* is clearly not normally distributed. Therefore, in principle, one may try to model the noise distribution of *Y* more precisely, and use a more targeted loss function than the *L*^2^-norm loss in (3). However, in our simulations we found that loss functions based on a binomial model for the pool sequencing procedure do not provide a significant improvement over the *L*^2^-norm loss and are computationally more challenging. This may be caused in part by the unpredictable variation of the bias term *B*. However, we added an option to our software that permits to replace (3) by weighted least squares in order to take differences in the coverage into account. That is, one can weigh the residuals according to the coverage depth. We found that this can improve the overall reconstruction in cases where coverage depth is very heterogeneous, as less weight is given to low-coverage and hence, high-variance, SNPs. We used this option on our mice example (see Section 4.2) since these data contain some sites with very low coverage.

Due to the discrete nature of *S*, the optimization problem in (3) is highly non-convex, which makes it challenging. However, conditioned on either of (*W, b*) or *S*, optimization in (3) becomes tractable: Indeed, minimizing (3) given (*W, b*) corresponds to a simple clustering problem with known centers

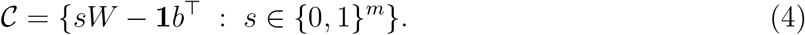

Given *S*, on the other hand, minimizing (3) corresponds to a simple linear regression problem with linear constraints on *W, b*. Thus, a very natural approach to tackle the minimization problem in (3) is to employ an iterative Lloyd’s type algorithm ^1^, see, e.g., [Lu and Zhou, 2016]. That is, to initialize either *S* or (*W, b*) and update iteratively until convergence of 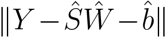.

Recently, [Lu and Zhou, 2016] showed that for a sub-Gaussian error distribution and appropriate initialization of either labels or clusters, Lloyd’s algorithm converges to an exponentially small clustering error in log(*N*) iterations. For generic Lloyd’s they also show that spectral clustering provides an appropriate initialization. Here, however, we cannot initialize the centers directly, but rather have to initialize the frequencies *W* and the bias term *b*, which indirectly determine centers via (4). For this, we propose to first cluster the *N* row vectors of *Y* ∈ [0, 1]^*N*×*T*^ into 2^*m*^ groups and then apply the HaploSepCombiExact (Algorithm 1 in the SI) to the 2^*m*^ estimated cluster centers 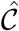. We summarize this initialization algorithm, of row clustering combined with the `HaploSepCombiExact`, in Algorithm 2 in the SI.

Recall that a combinatorial reconstruction as in the `HaploSepCombiExact` (Algorithm 1 in the SI) exactly recovers *W, S* and *b*, if *F* = *SW* + **1***b*^T^ is known exactly. Similarly, it can be shown, see [Behr and Munk, 2020], that the `HaploSepCombi` (Algorithm 2 in the SI) will recover *S* exactly and *W* (and *b*) up to some small error term, if the estimated cluster centers 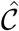 are sufficiently close to the true centers 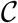 (= the row vectors of *F*) in (4). More precisely, this means that whenever 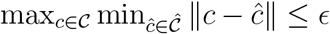, then it follows that 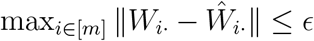, for any sufficiently small *E*, see [Behr and Munk, 2020] for details.

The pseudo-code in Algorithm 3 in the SI summarizes our complete procedure for iteratively recovering the haplotype structure *S* and frequencies *W* from data *Y*, using HaploSepCombi (Algorithm 2 in the SI) as initialization. The only tuning parameter of our procedure in Algorithm 3 is the threshold *δ* used in our iteration stopping criterion. We found that haploSep’s reconstruction is generally very robust w.r.t. *δ* and usually converges within a couple of iterations, see Section S2-3 in the SI for details. For our simulation results, as well as in our online R package implementation, we used *δ* = 0.001 as a default choice.

Note that in practice, the number of major haplotypes *m* is not given and has to be estimated from the data *Y*. Cross validation and bootstrap methods could be employed for this purpose. In Section S2-5, we present a different approach which is (computationally) much simpler and based on singular value decomposition. As further information on the reliability of our estimates, we propose accuracy measures and explain their computation in Section S2-5.

## 3 Simulations

Our simulations are designed to mimic experimental evolution (see, e.g.,[Kawecki et al., 2012], [Long et al., 2015] and [Schlötterer et al., 2015] for reviews). These experiments permit to study evolutionary adaptation under controlled laboratory conditions, making it easier to disentangle adaptive responses from other factors such as demography or genetic drift. Typically multiple populations of organisms are kept in the laboratory for several generations under stressful conditions chosen by the experimenter. DNA sequence information is commonly obtained at different time points to study the genetic basis of adaptive responses. However the separate sequencing of individuals at high coverage will often be too time consuming or costly for larger populations. As a consequence, the analysis is frequently carried out based on estimated population allele frequencies from pools of individuals sequenced together, although haplotype information would be helpful for a better understanding of the adaptive process.

With our simulations, we intend to illustrate how haplotypes and their relative frequencies can be reconstructed in such experiments. Furthermore, we show that it is possible to obtain improved population allele frequency estimates with pool sequencing experiments by using the reconstructed haplotypes and their estimated frequencies.

### 3.1 Reconstruction of haplotype structure and frequency

To illustrate our method, we discuss a viability selection model with a selected locus (selection coefficient *s* = 0.05) occurring on three of the haplotypes. Drift is simulated every generation via multinomial sampling from the haplotypes present in the previous generation. We consider an experiment with a constant population size over 150 generations. To mimic pool sequencing, the allele frequency data are obtained via binomial sampling at a Poisson (*λ* = 80) coverage from the population allele frequencies. We chose starting haplotypes from a set of founder haplotypes sequenced by [Barghi et al., 2019]. Due to the shared genealogy, such haplotypes share a large amount of similarity, making the reconstruction challenging. In this context, we consider three scenarios that differ with respect to the population size and the number of starting haplotypes. These parameters were chosen to mimic the experiments described in Section 4, namely, experiments with *C. elegans*, *D. simulans*, and the Longshank mice experiment. Further details concerning the simulation setup can be found in Section S3 (SI).

To better understand the performance of our estimates, some exemplary situations are shown in Fig. 1 and the respective allele composition results in Fig. S4 (SI). Whereas the allelic composition of the major haplotypes is estimated with very low error in Fig. 1(a), the haplotype frequency estimates become accurate only at later stages of this experiment. In Fig. 1(b), on the other hand, the frequency estimates are very accurate except at the very beginning of the experiment, but only the composition of the dominating haplotype is accurately estimated. Finally, in Fig. 1(c) both frequencies and allelic composition of the dominating haplotypes are estimated very accurately. The explanation for these observed patterns is that several low frequency confounder haplotypes are present for a considerable time in the Longshank mice experiment, before most of them get eliminated by genetic drift. With our *C. elegans* example on the other hand, genetic drift eliminates all except one haplotype very quickly. The remaining haplotype is easily estimated, but for the disappearing ones, only very few time points provide information to reconstruct their composition. Therefore the reconstruction error is between 24% and 37%. Finally for *D. simulans*, there is sufficient information in the data to estimate the two dominating haplotypes more or less perfectly. Frequency and allelic composition is accurately inferred even for a third one.

**Figure 1:**
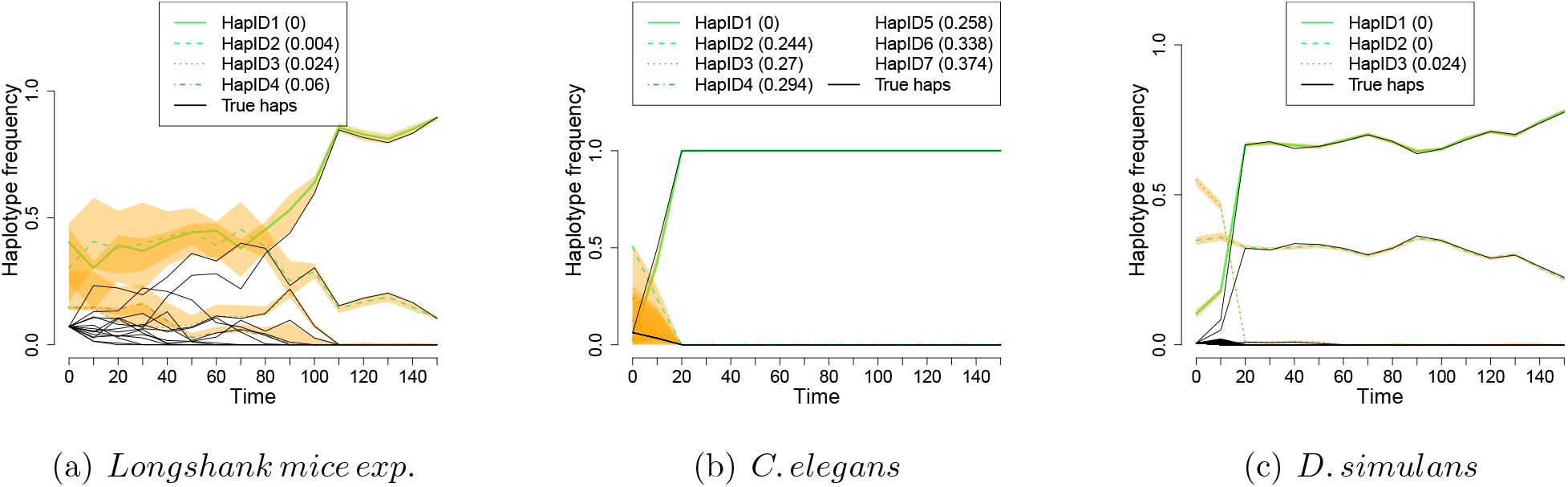
Reconstruction results. Results of a typical simulation run using features of (a) the Longshank mice experiment, (b) the *C. elegans* design, and (c) the *D. simulans* experiment. True (black solid lines) and reconstructed haplotype frequencies are shown, with accuracy intervals (0.025 to 0.975 quantiles based on bootstrap) in yellow. In parentheses, we report the proportion of mismatches between estimated and the corresponding true haplotype. Details about the simulation scenarios can be found in the text and Section S3 (SI).

For a more complete picture, we now report reconstruction errors for 100 simulation runs under the scenario mimicking the Longshank mice experiment in Fig. 2. The number of reconstructed haplotypes is estimated for each run via our model selection criterion explained in S2-4 (SI). The boxplots in Fig. 2(a) depict the errors in terms of the mismatch proportion for each of the reconstructed haplotypes, whereas Fig. 2(b) provides errors in terms of the mean absolute difference between the true and estimated frequencies at each time point. (See Section S1 for definitions of accuracies and box-plots). According to Fig. 2(a), the composition of the dominating haplotype (the one with the highest inferred frequency at the end of the experiment) is always estimated nearly perfectly. For the other haplotypes, the accuracy depends on whether the simulated trajectory provides enough information. As we simulated three of the haplotypes as selected, those haplotypes often (but not always) reached sufficiently high allele frequencies during the experiment and could therefore be estimated reliably. The frequency estimates in Fig. 2(b) clearly improve over time, illustrating again that accurate frequency estimates can be expected at time points where not too many haplotypes are present. Our simulations led to occasional outliers, i.e. situations where the accuracy is less satisfactory. For a practical application, we therefore recommend to use the accuracy scores *R*^2^ proposed in Section S2-5 and the frequency change of reconstructed haplotypes for assessing the reliability of our estimates. In Fig. 2 for instance, we filtered out scenarios where either *R*^2^ < 0.8 or the frequency change of dominating haplotype (HapID1) is below 0.1. These criteria generate a reasonable threshold to filter out problematic scenarios. For different experimental designs, we recommend to validate the thresholds with simulations. In supplementary Section S7, we provide a more detailed discussion of situations that may lead to outlying estimates. For results simulated under the other two experimental setups, see Figs. S5 and S6 (SI).

**Figure 2:**
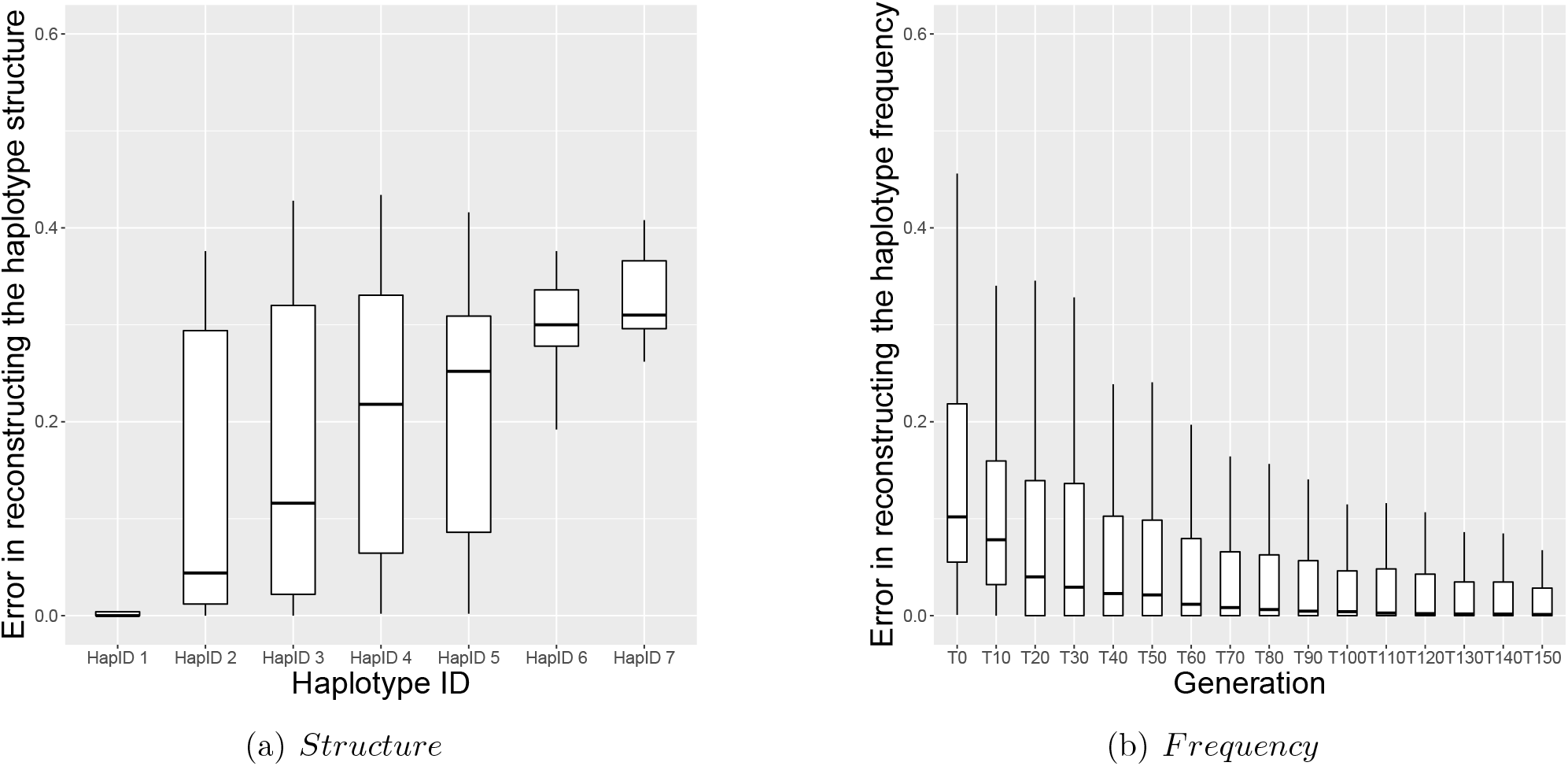
Haplotype reconstruction error. We consider our basic selection scenario with Longshank mice based on 100 simulation runs. (a) Proportion of wrongly classified SNPs for each reconstructed haplotype. The haplotypes are displayed in decreasing order according to their cumulative frequency over time. (b) Absolute difference between the true and estimated haplotype frequencies for each time point at which sequencing information is available. Box-plots definition can be found in Section S1 in the SI. As can be seen, error in reconstruction is low especially for the most abundant haplotype (hapID 1) and the late time points.

When planning an experiment, it can be useful to know under which design parameters haplotype reconstruction will tend to be reliable. For this purpose, we provide simulation results exploring the influence of E&R designs on the accuracy of our method and summarize the results in the supplement. See Section S3 for a summary of the simulated scenarios, and Section S4 for the results obtained under these scenarios. While our simulations suggest a good performance over a wide range of scenarios, we recommend that potential users perform additional simulations, if their experimental design deviates from our considered scenarios.

A general observation is that the more the haplotypes change in their frequencies during the experiment, the better the haplotype reconstruction gets. In E&R this can be achieved through a large enough selection pressure affecting the investigated genomic region, or through small population sizes such that genetic drift causes large frequency changes. In other applications, where samples may differ in location rather than time, a sufficient amount of population structure would be needed.

### 3.2 Improved allele frequency estimates

With known founder haplotypes, it has been shown in [Tilk et al., 2019] that allele frequency estimates from pool sequencing can often be improved by using haplotype information. Here we investigate, whether this observation also applies to the case of unknown founder haplotypes when using our estimates of important underlying haplotypes and their frequencies. Indeed, allele frequency estimates can be obtained by multiplying the matrix of the reconstructed haplotype structure 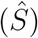 with the matrix of the estimated haplotype frequencies 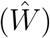 and adding the estimated bias term 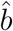. Using the simulated data in Section 3.1, we compared the so obtained estimates with the original allele frequencies from pool sequencing. As a measure of the difference in accuracy, we computed the ratio

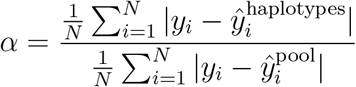

where *N* is the number of SNPs, *y*_*i*_ is the true allele frequency of SNP 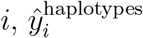 is the allele frequency of SNP *i* estimated using the reconstructed haplotypes, and 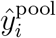 is the one estimated by pool sequencing. If *α* is smaller than one, the haplotype based estimate performs better. For each time point, we computed *α* based on all SNPs where the allele is not fixed or lost. To eliminate situations where the haplotype reconstruction does not work so well, we filter using the criterion in S2-5 to decide whether to use the haplotype based allele frequency estimates. Fig. 3 summarizes the relative performance for the Longshank mice experiment based on the filtered data. Analogous results for the other two experimental designs can be found in Fig. S17 (SI). Our results reveal that the use of the reconstructed haplotype information typically leads to an improved accuracy. Indeed, since haplotype frequency estimates combine information across many SNPs, they are less noisy than allele frequencies from pool sequencing for individual SNPs.

**Figure 3:**
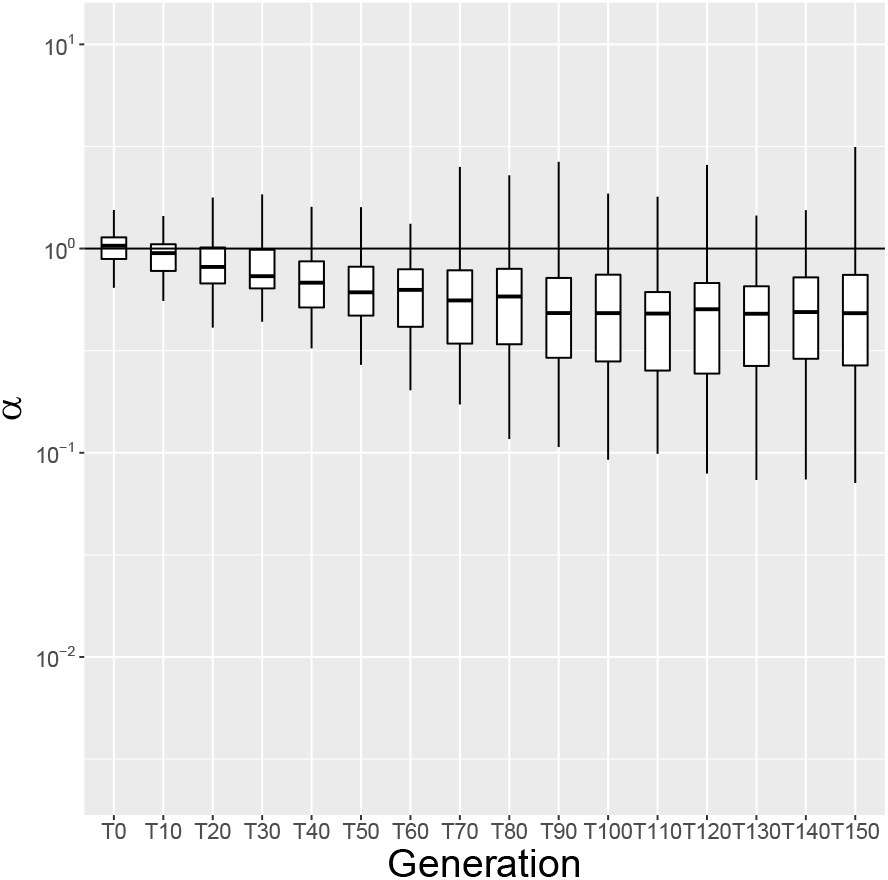
Comparison between pool sequencing and haplotype reconstruction estimates of the allele frequencies. Error ratio (*α*) between haplotype based allele frequency estimates (numerator) and the pool sequencing estimates (denominator) plotted on a log-scale. Results from 100 simulations based on the Longshank mice experimental design. Except for the first time point (*T*0), the haplotype based estimates are usually more accurate.

### 3.3 Recombination

Recombination is an evolutionary force that occurs with sexual reproduction and exchanges genetic material between chromosomes. In higher organisms, it constantly produces new haplotypes, so that none of the chromosomes will exist in its original form after some generations. Since our method requires the reconstructed haplotypes to be present in the same form for a sufficient number of samples, we applied it to DNA segments short enough to make recombination an infrequent event. We stress that shorter window sizes are often of particular interest, for example, in E&R experiments, where researchers want to understand the haplotype dynamics underlying fairly narrow selected blocks. (E.g., in [Barghi et al., 2019], selected block lengths between 1.65 kb and 5 Mb were observed.) Due to different SNP densities and recombination rates, an appropriate segment size for our method will differ between experiments and organisms.

The recombination rate *r* is commonly measured in centiMorgans per megabase (cM/mb), with 1 cM/mb representing an expected number of 0.01 cross-over events on a sequence piece consisting of 10^6^ bases. In our simulations, we focused on a genomic region of length 10000bp containing 500 linked SNPs which leads to a realistic SNP density for *D. simulans* [Barghi et al., 2019]. According to [Howie et al., 2019], *r* is approximately 3.4 cM/mb for this species, so that recombination will occur on average only in 3.4 out of 10000 meioses. For mice, *r* ≈ 0.528 cM/Mb on average, but short hotspot regions are immersed within a low recombining background. Recombination for *C. elegans* varies greatly in the genome and across experiments as selfing can be enforced as well. The average recombination rate has been estimated as 3.06 cM/mb in [Barnes et al., 1995]. The SNP density for our considered region in the mice data leads to a region of approximately 110000bp for 500 SNPs, whereas for the *C. elegans* experiment 500 SNPs would correspond to a region of approximately 32000bp. Overall, these quantities suggest that recombination plays only a minor role in our analysis involving the above organisms.

To better understand the effect of recombination on our reconstruction, however, we also explored situations where *r* is much larger. According to our simulations explained in Section S8 in the SI, even a more than five times higher recombination rate than for *Drosophila* left the accuracy of the reconstructed haplotypes almost unchanged. (See Figure S23). This is interesting, since the number of haplotypes present in the samples increased a lot with high recombination rates (see Fig. S21). A reason for this result could be that only a very small proportion of recombination products will reach high enough frequencies in order to get reconstructed by our method. Indeed, Fig. S22a suggests that high recombination rates produce a large fraction of low frequency haplotypes that make the haplotype frequency estimates slightly less accurate (Fig. S22b). On the other hand, the proportion of simulation runs that passed our accuracy score threshold, and was therefore considered sufficiently reliable for usage, decreased with increasing recombination rate *r.* Overall these results suggest that haploSep works at least in some cases also under considerable recombination which would make reconstruction feasible also with genomic segments longer than those considered in our examples. In future work we plan to investigate approaches to extend our haplotype reconstruction to even larger regions.

## 4 Application to real data

We applied our approach to the E&R data sets taken from [Barghi et al., 2019], and [Noble et al., 2019], as well as to the HIV longitudinal data from [Zanini et al., 2015], and to the not yet published data set described in [Castro et al., 2019]. In each case, we looked at a small genomic region and applied our method there. Using the data sets from [Barghi et al., 2019], [Noble et al., 2019] and [Zanini et al., 2015], we also compared our inferred haplotypes with reference haplotypes provided by the authors.

As a further validation of our approach, we compared our reconstructed haplotypes with paired end reads using the original sequencing data from [Barghi et al., 2019]. We found our reconstructed haplotypes to be vastly concordant with the reads when interpreting them as very short haplotypes. For further details see Section S9 (SI).

### 4.1 Drosophila simulans (D. simulans)

We now consider the E&R experiment of [Barghi et al., 2019]. There the base population consists of 202 isofemale lines of *D. simulans*. Ten replicate populations were kept for 60 generations, with sequencing data available every ten generations. Furthermore, a sample of 189 founder haplotypes was sequenced, as well as 100 additional ones from five evolved replicates.

For this data, we first identified interesting genomic regions by testing for signals of selection at the SNP level using the modified *χ*^2^ and Cochran–Mantel–Haenszel tests [Spitzer et al., 2020] that account for drift and sequencing noise in the data. We then applied our method to multiple regions that show statistically significant allele frequency changes. In particular, we considered positions 11.239636 to 11.591566 Mb on chromosome 2L for replicate 3. Fig. 4 provides the estimated haplotype trajectories, as well as a comparison between our estimated allelic composition and the best matching founder haplotype sequences (see Section S1 in the SI for a definition of best matching haplotypes). Due to the presence of a large number of similar haplotypes in particular at the early generations, the reconstruction is quite challenging for this experiment. Nevertheless, the dominating haplotype is usually reconstructed almost without error.

**Figure 4:**
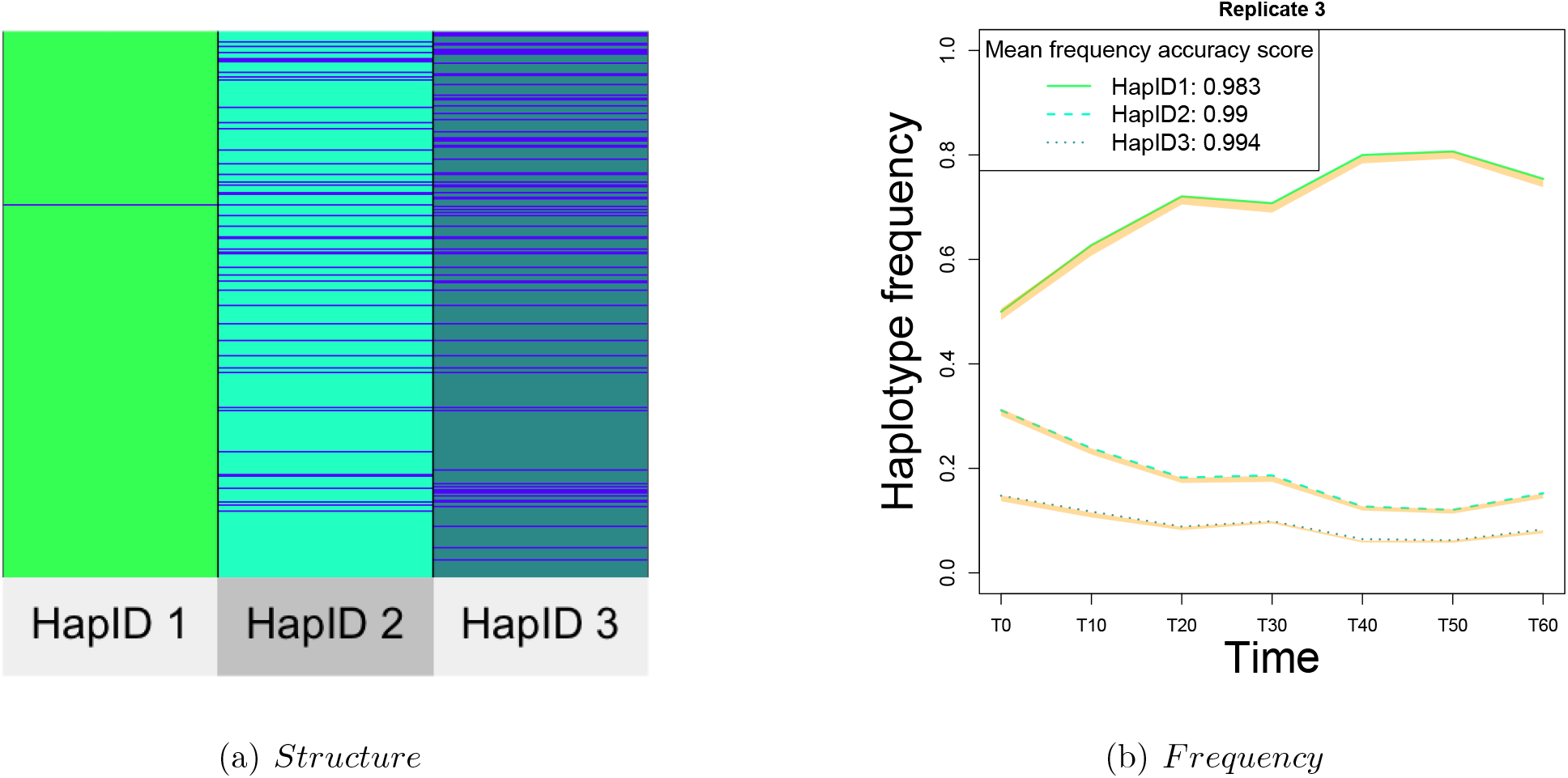
Reconstruction results for an example region from [Barghi et al., 2019]. (a) Match between reconstructed haplotype structure and sequenced founder haplotypes for the region 11.239636 to 11.591566 Mb on chromosome 2L (replicate population 3) from the *D. simulans* experiment. Blue lines indicate mismatches. This plot is generated using the superheat function from [Barter and Yu, 2018]. (b) Reconstructed haplotype frequencies with accuracy intervals (in yellow) and mean accuracy scores, with data as in (a). See details on the accuracy intervals in Section S2-5 and S1.

### 4.2 Longshank experiment in mice

In the Longshank mice experiment, individuals from a mouse population were selected to produce offspring according to their tibia length/body mass ratio. The evolution of three populations (two Longshank lines and one control line) was followed over several generations. For details on the experimental design, we refer to [Castro et al., 2019] and [Marchini et al., 2014]. We received time series data from this experiment for the the Nkx3-2 region (4395 SNPs and indels) of the Longshank 1 (LS1) line collected every generation, from generation 0 to 20 (still unpublished). The allele frequencies were missing for a large number of SNPs. Therefore we decided to remove the later generations (14-20) from our analysis, because of their particularly high proportion of missing values. For the generations 0-13, we only kept those SNPs for which all allele frequencies were available. Filtering the data this way, we reconstructed haplotypes from the remaining 561 SNPs in this region.

The right panel of Fig. S26 displays our estimated haplotype trajectories and the corresponding accuracy scores. Unfortunately no founder haplotypes or read data for comparison purposes were available to us with this experiment.

### 4.3 Caenorhabditis elegans (C. elegans)

We next look at the experiment described in [Noble et al., 2019]. There, three replicate populations experienced an increasing quantity of NaCl during their evolution. The base population comprised 10^4^ individuals that originated from 16 founder inbred lines. Pool sequenced allele frequency data have been made available to us for the base population and for the evolved populations at generations 50 and 100. Additionally, sequence information for the 16 founder inbred lines, as well as low coverage sequencing of a few individuals from the base population and the two subsequent time points has been provided. Further details on the experimental design can be found in [Noble et al., 2017].

As for section 4.1 we searched for genomic regions showing signatures of selection. As an example, we provide results for a genomic region containing 666 SNPs (chromosome 5, 14924777-15216613 bp). The upper panel of Fig. S27 (SI) illustrates the close match between the reconstructed haplotypes and the most similar sequenced founder haplotype. For each replicate line, the lower panel of Fig. S27 (SI) shows the reconstructed haplotype trajectories, together with the corresponding accuracy measures. Moreover, since the three replicates show similar evolutionary patterns, we decided to also apply our method to all of them simultaneously. The result seems consistent with our single replicate analysis in terms of the reconstructed haplotypes and their frequency trajectories. This suggests parallel evolution across the replicates (see Fig. S28 in the SI).

### 4.4 HIV

To further illustrate the broad applicability of our method, we now present an example involving HIV. We used data from the first patient in the longitudinal data set of [Zanini et al., 2015], available at https://hiv.biozentrum.unibas.ch/data/ (downloaded 19/11/2020). Twelve samples are available for the chosen patient who got infected with HIV-1 of type AE. The samples were obtained within a time window from 122 to 2996 days after infection. HIV has an extremely high mutation rate (4.1 × 10^−3^ per bp, compared to 4.65 × 10^−9^ for *Drosophila*). Therefore new SNPs are constantly introduced. Although `haploSep` has been designed to reconstruct stable haplotype structures, it can still be used to estimate dominant haplotype structures that do not change much over time. Given the relatively short genome (9060 base pairs), we decided to reconstruct haplotypes on a genome wide scale. Using the model selection step discussed in Section S2-4 (SI), four haplotypes were obtained. From the reconstructed haplotype information, we computed allele frequency estimates as described in Section 3.2. As shown in Fig. S29c (SI) the observed allele frequency dynamics is well represented using only four haplotypes. Pieces of the true genomes have been made available by [Zanini et al., 2015]. Therefore, we were able to compare true and reconstructed haplotypes on a 245 bp window with 53 SNPs. Although the true haplotypes change between time points due to mutations, the median mismatch to our estimates is only 5.7% (see Fig. S29a in the SI). We refer to Fig. S29b (SI) for frequency estimates and to Section S13 (SI) for a comparison with another method.

## 5 Method Comparison

Methods have been proposed for reconstructing haplotype information from pool sequence data that use approaches completely different from ours. Most of these methods use the linkage information contained in read data. To our knowledge, read based methods have been developed with genomes from viruses and microorganisms as the main application. Due to the massive computational effort required, the short genomes in this context make such an approach computationally feasible. For other model organisms, as considered in this work, these read based methods can be computationally extremely challenging, and we were not able to run them on the examples considered in this work. Therefore, we focused on scenarios that mimic HIV evolution and involve much fewer SNPs. We chose `CliqueSNV` [Knyazev et al., 2018] for our comparison, because it has been developed recently and ran on most of these simplified scenarios. Although, `CliqueSNV` is much slower than `haploSep`, it is still fast compared to other read based methods.

With the exception of [Franssen et al., 2017], methods that use only allele frequency information require the haplotype sequences to be known in advance (e.g. [Long et al., 2011], [Kessner et al., 2013] and [Cao and Sun, 2015]). Estimating only the haplotype frequencies is a much simpler task than reconstructing both structure and frequency as achieved by `haploSep`. The heuristic method of [Franssen et al., 2017] works only on windows in the genome that have been affected by selection. On such a window, only one haplotype is reconstructed for a small subset of SNPs that exhibit highly correlated allele frequency trajectories across multiple time series. Our approach, on the other hand, does not require replicated experiments and is applicable under much more general circumstances. Due to the different haplotype concept used, it is not directly comparable with [Franssen et al., 2017].

Although our method was not designed for haplotype dynamics in viruses (see Section 4.4), we compared `haploSep` with `CliqueSNV` (version 1.5.3, downloaded 12/11/2020) on simulated data and the real HIV data from [Zanini et al., 2015]. Here we present the major findings from our comparison. We first follow the HIV simulation scenario suggested in [Cao et al., 2020]. We refer to SI Section S13-1 for details. Due to the long run times for data generation and the analysis with `CliqueSNV`, we only carried out 20 simulations each involving 24 samples. Unfortunately, six simulated data sets contained samples where `CliqueSNV` crashed, whereas `haploSep` provided results for all samples. For our comparison in Fig. 5 we used all samples for which results could be obtained, leading to 480 samples for `haploSep` and 470 for `CliqueSNV`. According to Fig. 5, `haploSep` provides better frequency estimates, but slightly less accurate structure estimates. This may be because `haploSep` provides identical reconstructions across samples while the true haplotypes change due to the high mutation rate used in the simulations. For a more detailed comparison of both methods see the error quantiles provided in Section S13. In terms of the allele frequency reconstruction (see Section 3.2), both methods show a very similar performance according to Fig. S30.

**Figure 5:**
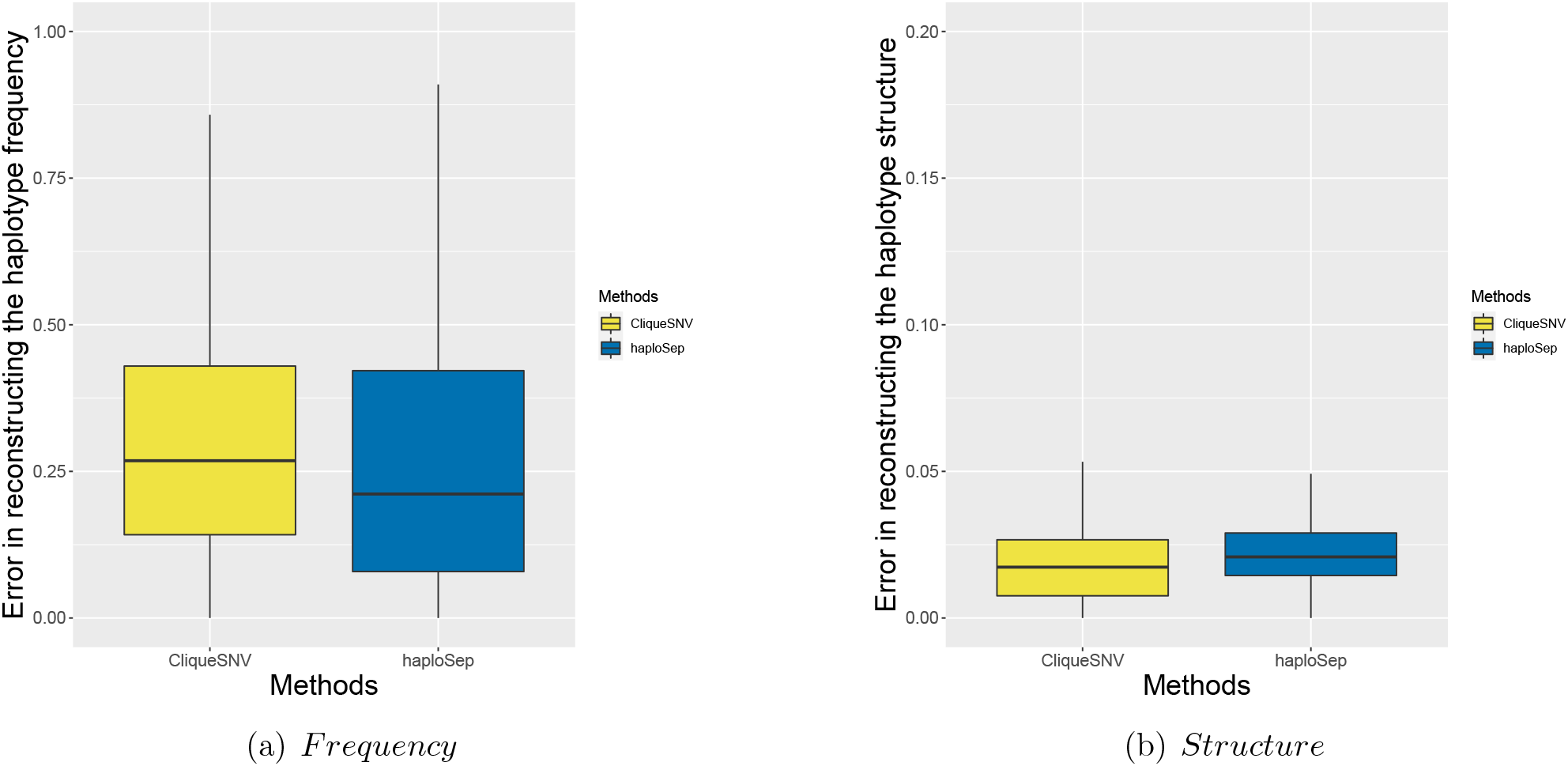
Comparison of `haploSep` and `CliqueSNV`. This figure considers all available samples and haplotypes from the 20 simulated data sets. For the structure, reconstruction errors were computed as proportion of mismatches between true and estimated haplotype structure among the SNP positions where the observed allele frequency is larger than 0.05 in all samples. For the frequencies, the absolute value of the difference between estimate and truth is used. See Section S1 for details on the error computations. (a) Errors of the haplotype frequency estimates. (b) Errors of the haplotype structure estimates. As can be seen, the accuracy of `haploSep` is higher for frequency estimates, but slightly lower for structure estimates.

It seems worth mentioning that our proposed approach took 12–94s depending on the simulation run, which is much faster than `CliqueSNV` (run times 900–18,000s). As the running time of `CliqueSNV` increases quadratically with the number of SNPs, compared to linear for `haploSep`, we would expect much larger differences in run time for regions containing more than the 200 SNPs used in our example.

For the real HIV data from [Zanini et al., 2015] our comparison focuses on the same 245bp window as in Section 4.4. Although recombination can occur with retroviruses such as HIV, mostly mutation leads to rapid temporal changes in our haplotype sequences. Due to this lack of stability, we compare our estimates with the respective closest true haplotype at each considered time point. As mentioned in Section 4.4, the availability of short segments of the real haplotypes and their frequencies makes a comparison with the truth possible. According to Fig. S33 in the SI, `haploSep` estimates the structure more accurately. On the other hand the frequencies are inferred more precisely by `CliqueSNV` (see Fig. S34 in the SI).

We also tried to compare `haploSep` and `CliqueSNV` on a small segment of 95 SNPs from the data set in [Barghi et al., 2019]. Unfortunately `CliqueSNV` crashed on our Mac Pro machine with 2,7 GHz 12-Core Intel Xeon E5 Processor with 32 GB RAM due to memory errors. We therefore compared `haploSep` with another method by [Cao et al., 2020] that took 2631 minutes (≈158000*s*) to run. See SI Section S13-4 for details.

In summary, given the high mutation rate, our method performs remarkably well in terms of accuracy both on the real and simulated HIV examples without requiring additional linkage information from reads. A major advantage of our method is its speed and computational stability. Compared to read based methods, it does not require a lot of memory and runs therefore on much longer sequences.

## 6 Discussion

We proposed a new, computationally efficient principled approach that estimates for the first time multiple completely unknown haplotypes from allele frequency data only. Under a suitably chosen experimental design, and with sufficiently large allele frequency changes, the allelic composition of the unknown haplotypes can be recovered reliably. Strong enough selection provides one scenario that leads to sufficient fluctuations in allele frequency.

Our method `haploSep` builds on a computationally efficient Lloyd’s type algorithm, which iteratively updates the discrete SNP structure and the respective frequencies of dominant haplotypes in a population. By efficiently exploring the combinatorial structure induced by the discrete haplotypes, we obtain an accurate initialization for the iteration. This enables `haploSep` to reconstruct several haplotypes containing thousands of variable positions from multiple samples within a couple of seconds on a standard laptop, while requiring bulk allele frequency data only. Existing methods that work on read data require much longer run times (see Section S13 in the SI).

A good reconstruction of haplotype frequencies is achieved at time points when a moderate number of haplotypes is present at a sufficiently high frequency which may not be the case early on when an experiment starts with many founder haplotypes. Further important design parameters that affect the quality of our reconstruction are the population size, and the number of samples with sequence information.

A lot of scientific studies use haplotype information to answer their research questions. By providing estimates of the most important underlying haplotypes, our method will help researchers that only have allele frequency data available.

Our estimated haplotype frequencies can also be used to obtain allele frequency estimates that are less noisy on average than the original frequency data obtained for instance via pool sequencing. Indeed, by combining information from several neighboring SNPs, the sampling variation introduced by sequencing a whole pool of individuals gets averaged out to some extent.

As our next step, we plan to extend our approach to data from locally structured populations, where samples are usually taken from multiple sub-populations. As another application, the reconstructed haplotypes may also be helpful to impute missing data from low coverage sequencing. In view of recombination, a challenging topic for subsequent research would be to extend our method to work on much longer genomic regions.

## Acknowledgement

We are grateful to the laboratories of Nick Barton, Christian Schlötterer, and Henrique Teotonio for providing us with their experimental data. We would also like to thank Quan Long, Lauren Mak, and Chen Cao, for helping us in using PoolHapX. This work has been supported by the Austrian Science Fund (FWF Doctoral Program Vienna Graduate School of Population Genetics”, DK W1225-B20). MB was supported by Deutsche Forschungsgemeinschaft (DFG; German Research Foundation) Postdoctoral Fellowship BE 6805/1-1. Moreover, MB acknowledges funding of DFG-GRK 2088. This work benefited from a research stay that was partially supported by the Simons Foundation and by the Mathematisches Forschungsinstitut Oberwolfach. AM and MB acknowledge support of DFG-SFB 803 Z02. AM and HL are funded by the Deutsche Forschungsgemeinschaft (DFG, German Research Foundation) under Germany’s Excellence Strategy – EXC 2067/1-390729940.

## Data and Code availability

All our code is written in R. Our software, simulation functions, and examples are available under https://github.com/MartaPelizzola/haploSep. We used published data from [Barghi et al., 2019], [Noble et al., 2019], and time series data from the experiment described in [Castro et al., 2019] (only partially published so far).

## Author contribution

AF and AM conceived the project. MP, MB, HL, and AF contributed to the design of the research and wrote the manuscript. MP, MB, and HL wrote the code for software, simulations and data analysis. All authors read and approved the manuscript.

## Footnotes

1 Instead of a hard iterative update of *S* and (*W, b*) as in a Lloyd’s type algorithm, one could also consider an EM-type algorithm, which updates the full distribution of the discrete matrix *S* ∈ {0, 1}^*N*×*m*^, whose *N* rows determine to the membership to the 2^*m*^ cluster centers of the rows of *Y*. However, it is not clear how such an approach can be implemented in a computationally efficient way. The reason for this is that in our case each of the *N* different rows of *Y* potentially depends on the full matrix *W*. Hence, in the maximization step, one would need to average over all 2^*N*×*m*^ possible binary matrices in {0, 1}^*N*×*m*^ for *S*, which is computationally not feasible, due to the exponential dependence on *N* (with *N* typically larger than 100).

