## Supplementary Material for "Multiple Haplotype Reconstruction from Allele Frequency Data"

Marta Pelizzola[1,2,a], Merle Behr[3,a], Housen Li[4], Axel Munk[4,5], Andreas Futschik[6, b]

1 Vetmeduni Vienna
2 Vienna Graduate School of Population Genetics
3 University of California Berkeley
4 University of Göttingen
5 Max Planck Institute for Biophysical Chemistry, Göttingen
6 Johannes Kepler University Linz
a These authors contributed equally

Our supplementary material is structured as follows. We first provide additional information on our proposed method in Section S2. In particular, we discuss conditions that ensure identifiability, i.e. unique estimates for our underlying haplotypes and their frequencies. We also provide algorithms and explain how we select the number of haplotypes (model selection), and how accuracy scores are computed that provide information on the quality of the estimates.

In Section S3, we describe our model for the simulations. We provide additional results from our simulations, together with our analysis of the error under several experimental designs, in Section S4. We evaluate the accuracy measure introduced in Section S2-5 with our simulations in Section S5. Furthermore, additional results on the estimation of allele frequencies are provided in Section S6. Section S7 provide an analysis of the simulation runs leading to outliers in the reconstruction error and Section S8 discusses the effects of different levels of recombination on our proposed approach. Additional results on the real data can be found in Sections S9, S10, and S11, S12. Lastly, Section S13 presents further details and results on the comparison with other methods.

### S1 Definitions and Notation

- **Box-plots:** For boxplots, the lower and upper hinges correspond to the first and third quartiles. The whiskers extend to 1.5 × IQR from the hinges (where IQR is the interquartile range). The same applies to all boxplots in the manuscript.
- **Haplotype structure accuracy:** The accuracy for the haplotype structure is computed as the proportion of mismatches between true and estimated haplotype structures.
- **Haplotype frequency accuracy:** The accuracy for the haplotype frequency is computed as the absolute value of the difference between true and estimated frequency for each haplotype at each time point.
- **Allele frequency accuracy:** The accuracy in allele frequency ($\alpha$) is computed per sample as $\alpha = \frac{\frac{1}{N}\sum_{i=1}^{N}|y_i - \hat{y}_i^{\text{haplotypes}}|}{\frac{1}{N}\sum_{i=1}^{N}|y_i - \hat{y}_i^{\text{pool}}|}$ where $N$ is the number of SNPs, $y_i$ is the true allele frequency of SNP $i$, $\hat{y}_i^{\text{haplotypes}}$ is the allele frequency of SNP $i$ estimated using the reconstructed haplotypes, and $\hat{y}_i^{\text{pool}}$ is the one estimated by pool sequencing.
- **Most frequent haplotype:** Haplotype having the highest sum of the frequency over all samples.

- **Best matching haplotype:** True haplotype having the most similar structure to a given reconstructed haplotype.

- **Haplotype frequency accuracy intervals:** The accuracy intervals for haplotype frequencies are the 0.025 and 0.975 quantiles of $\hat{W}_{it}(Y^*)$ as detailed in S2-5.

### S2 Theory and Methods

#### S2-1 Identifiability of structure and frequency from allele frequency (AF)

[Behr and Munk, 2017] derived sufficient and necessary conditions under which the matrices $S$ and $W$ (including the number of haplotypes $m$) are uniquely identifiable from their product $SW$. With some slight modifications of their arguments, we can also show that under weak identifiability assumptions on $S$ and $W$, one can uniquely identify $S$, $W$, and $b$ from the population AFs $F$. More precisely, for $W$ it is assumed that different combinations of SNPs lead to different AFs, that is,

$$sW \neq s'W \text{ for all } s \neq s' \in \{0,1\}^m. \tag{S1}$$

For the haplotype structure $S$ it is assumed that there is at least one SNP which is unique to a haplotype and at least one SNP that is only present in minor haplotypes, that is

$$\begin{aligned} &\text{for all } i \in [m] \text{ there exists an } n \in [N] \text{ such that} \\ &\qquad S_{ni} = 1 \text{ and } S_{nj} = 0 \text{ for all } j \neq i \\ &\text{and there exists an } n \in [N] \text{ such that } S_{ni} = 0 \text{ for all } i \in [m], \end{aligned} \tag{S2}$$

(equivalently one can exchange 0 and 1 in (S2)). Both of these conditions are very reasonable in most real data situations, given that the number of essential haplotypes $m$ is not too large. It is easy to see that condition S1 is necessary for identifiablity of haplotype structure $S$ and frequency $W$ from AF $Y$ in (2). A simple situation, where S1 does not hold is when two haplotypes have exactly the same proportion at all time points $t \in [T]$. In that case, it is not possible to distinguish whether a SNP is present in one or the other haplotype. Condition (S2) imposes a sufficient variability of individual haplotypes. A trivial non-identifiable counter example is, for instance, when one major haplotype is constant zero or constant one. Some further insights and examples on the specific condition in (S2) can be found in [Behr and Munk, 2017, Behr et al., 2018]. Note that (S2) requires that out of the $2^m$ possible variant combinations for the $m$ haplotypes, at least those $m$ combinations which correspond to the identity vectors $e_1 = (1, 0, \ldots, 0), \ldots, e_m = (0, \ldots, 0, 1)$ and the one which corresponds to the zero vector $(0, \ldots, 0)$ appear at some of the locations $n \in [N]$.

The conditions (S1) and (S2) do not just guarantee identifiability in an abstract way, but they also lead to an explicit algorithm for recovering $S$, $W$, $b$ and $m$ from the noiseless AFs $SW + b$ in (2). Part of our reconstruction algorithm is built on this deterministic recovery algorithm that is based on a simple combinatorial reordering of the observations (see [Behr and Munk, 2017, Diamantaras and Chassioti, 2000] for very similar algorithms). The idea of this algorithm is that the discrete nature of $S$ lets us identify both $S$ and $W$ from appropriate row vectors of $Y$ as outlined in the following.

The smallest norm among the rows of $Y$ appears for any SNP that has variant 0 for all $m$ haplotypes, in which case we observe only the bias term $b$. Similar, the second (and third) smallest possible row value of $Y$ appears for a SNP with variant 0 on all haplotypes, except the one with the smallest frequency $W_{m\cdot}$ (second smallest frequency $W_{(m-1)\cdot}$), which lets us identify $W_{m\cdot}$ and $W_{(m-1)\cdot}$. Among the remaining observed row values of $Y$ the smallest one must correspond to $W_{(m-2)\cdot}$, and so on. In that way, one can successively recover all the frequencies $W_{i\cdot}$ and given $W$ it is straightforward to recover $S$. We present pseudo code in Algorithm 1 below.

#### S2-2 Algorithms

**Algorithm 1** Recover $S, W, b$ from exact data $Y = SW + b$

**procedure** HAPLOSEPCOMBIEXACT

**Input:** $Y = SW + \mathbf{1}b^\top$ such that (S1) and (S2) hold.

**Output:** $S, W, b, m$

    $\mathcal{Y} \leftarrow \{Y_{1\cdot}, \dots, Y_{N\cdot}\}$
    $b \leftarrow \arg\min_{y \in \mathcal{Y}} \|y\|$
    $\mathcal{Y} \leftarrow \mathcal{Y} \setminus b$
    $\mathcal{Y} \leftarrow \mathcal{Y} - b$
    $W_{1\cdot} \leftarrow \arg\min_{y \in \mathcal{Y}} \|y\|$
    $\mathcal{Y} = \mathcal{Y} \setminus W_{1\cdot}$
    $m \leftarrow 1$
    **while** $\mathcal{Y} \neq \emptyset$ **do**
        $W_{(m+1)\cdot} \leftarrow \arg\min_{y \in \mathcal{Y}} \|y\|$
        $m \leftarrow m + 1$
        $\mathcal{Y} \leftarrow \mathcal{Y} \setminus \{\sum_{i=1}^{m} s_i W_{i\cdot} \; : \; s \in \{0,1\}^m\}$
    **end while**
    **for** $n = 1$ to $N$ **do**
        $S_{ni} \leftarrow \arg\min_{s \in \{0,1\}^m} \|Y_{n\cdot} - sW\|$
    **end for**
    put $W_{i\cdot}$ in the reverse order
    **return** $S, W, b, m$

**end procedure**

**Algorithm 2** Initialize $\hat{W}, \hat{b}$ from $Y$ in (2)

**procedure** HAPLOSEPCOMBI<br>
**Input:** $Y \in [0,1]^{N \times T}$ and $m \in [N]$<br>
**Output:** $\hat{W}, \hat{b}$<br>
    $\{C_1, \ldots, C_{2^m}\} \leftarrow$ hierarchical clustering of $\{Y_{n\cdot} : n \in [N]\}$ with $2^m$ centers $\subset [0,1]^T$.<br>
    $\hat{\mathcal{C}} \leftarrow \{C_1, \ldots, C_{2^m}\}$<br>
    $\hat{b} \leftarrow \arg\min_{c \in \hat{\mathcal{C}}} \|c\|$<br>
    $\hat{\mathcal{C}} \leftarrow \hat{\mathcal{C}} \setminus b$<br>
    $\hat{\mathcal{C}} \leftarrow \hat{\mathcal{C}} - b$<br>
    $\hat{W}_{1\cdot} \leftarrow \arg\min_{c \in \hat{\mathcal{C}}} \|c\|$<br>
    $\hat{\mathcal{C}} \leftarrow \hat{\mathcal{C}} \setminus \hat{W}_{1\cdot}$<br>
    **for** $l = 2$ to $m$ **do**<br>
        $\hat{W}_{l\cdot} \leftarrow \arg\min_{c \in \hat{\mathcal{C}}} \|c\|$<br>
        **for** $s \in \{0,1\}^{l-1}$ **do**<br>
            $\hat{\mathcal{C}} \leftarrow \hat{\mathcal{C}} \setminus \{\arg\min_{c \in \hat{\mathcal{C}}} \|c - \sum_{i=1}^{l-1} s_i \hat{W}_{i\cdot} - \hat{W}_{l\cdot}\|\}$<br>
        **end for**<br>
    **end for**<br>
    **return** $\hat{W}, \hat{b}$<br>
**end procedure**

**Algorithm 3** Recover $S, W, b$ from $Y$ in (2)

**procedure** HAPLOSEP<br>
**Input:** $Y \in [0,1]^{N \times T}, m \in [N], \delta > 0$<br>
**Output:** $\hat{W}, \hat{b}, \hat{S}$<br>
    $(\hat{W}, \hat{b}) \leftarrow$ HAPLOSEPCOMBI$(Y, m)$<br>
    **for** $n = 1$ to $N$ **do**<br>
        $\hat{S}_{n\cdot} \leftarrow \arg\min_{s \in \{0,1\}^m} \|Y_{n\cdot} - s\hat{W} - \mathbf{1}\hat{b}^\top\|$<br>
    **end for**<br>
    $E_0 \leftarrow 0$<br>
    $E_n \leftarrow \|Y - \hat{S}\hat{W} - \mathbf{1}\hat{b}^\top\|$<br>
    **while** $|E_n - E_0| > \delta$ **do**<br>
        $E_0 \leftarrow E_n$<br>
        $(\hat{W}, \hat{b}) \leftarrow \arg\min_{W,b} \|Y - \hat{S}W - \mathbf{1}b^\top\|$<br>
        such that $W_{it}, b_t \in [0,1]$, $\sum_{i=1}^m W_{it} \leq 1$<br>
        **for** $n = 1$ to $N$ **do**<br>
            $\hat{S}_{n\cdot} \leftarrow \arg\min_{s \in \{0,1\}^m} \|Y_{n\cdot} - s\hat{W} - \mathbf{1}\hat{b}^\top\|$<br>
        **end for**<br>
        $E_n \leftarrow \|Y - \hat{S}\hat{W} - \mathbf{1}\hat{b}^\top\|$<br>
    **end while**<br>
    **return** $\hat{W}, \hat{b}, \hat{S}$<br>
**end procedure**

#### S2-3 Computational aspects of haploSep

In the following we provide more details on computational aspects of the `haploSep` procedure. Recall that `haploSep` takes as input a matrix $Y \in [0,1]^{N\times T}$ with allele frequency data, as well as an integer $m$, which gives the number of estimated haploypes. From a computational perspective, the relevant regime for haplotype reconstruction is when $N$ is large (typically larger than 100), $T$ is of small or moderate size (typically smaller than 100) and $m$ is small (typically around $2-8$). In the following, we consider each of the different steps in the `haploSep` procedure separately and analyze computational aspects.

1. **(Clustering)** In the `haploSepCombi` initialization algorithm, see Algorithm 2, the first step is to cluster the $N$ rows of the matrix $Y$ into $2^m$ groups. To this end, we employed hierarchical clustering via the R function `hclust` from the R package `stats` with Eucledian distance metric. The complexity to compute the distance matrix between the $N$ different rows, each of dimension $T$, is $\mathcal{O}(N^2T)$. For the whole `haploSep` procedure, this is the only part which has a quadratic dependence on the number of sample $N$ (all other steps are linear in $N$) and hence, for a typical sample size regime in haplotype separation, this part is the computational bottleneck of the current implementation. Nevertheless, for all the scenarios considered in this paper, the overall computation time of `haploSep` never took longer than a few seconds on a standard laptop. If needed, however, one may replace hierarchical clustering with a computationally faster algorithm, as, e.g., k-means, in which case the overall computational complexity of `haploSep` will be linear in the number of variants $N$.

2. **(Combinatorial Initialization)** Given the $2^m$ cluster centers, each of dimension $T$, from the previous step, `haploSepCombi` as in Algorithm 2 then reconstructs an estimate for the haplotype frequency $\hat{W}$ and the bias term $\hat{b}$. Note that $2^m \ll N$. Thus, the computation time of this part is completely independent of $N$ and therefore typically negligible. More precisely, computation time of this part is of order $\mathcal{O}(2^{2m}T)$.

3. **(Lloyd's-type Iteration)** Given the initialization $(\hat{W}, \hat{b})$ from the previous step, `haploSep` (see Algorithm 3) then iteratively updates $(\hat{W}, \hat{b})$ and $\hat{S}$ until convergence. An individual update step of $\hat{S}$ amounts to comparing the distances of the $N$ rows of $Y$ to the current $2^m$ centers as in (4), each of dimension $T$, and thus, has computation time $\mathcal{O}(2^m NT)$. An individual update step of $(\hat{W}, \hat{b})$ amounts to linear regression (with convex constraints), which has a linear worst case computational complexity w.r.t. the number of sample $N$ (as well as a linear computational complexity w.r.t. the number of time points $T$). To solve this part efficiently, we use the `lsei` function form the R package `limSolve`. Note that both update steps of $\hat{S}$ and of $(\hat{W}, \hat{b})$ result in a monotone decrease of the overall $L^2$ error, $\|Y - \hat{S}\hat{W} - \mathbf{1}b^\top\| \geq 0$ and therefore, will converge eventually. In practice, we found that `haploSep` usually converges within a couple of iterations, and for any stopping threshold $\delta \leq 0.001$ (in our simulations we chose $\delta = 0.001$) results were almost completely independent of the choice of $\delta$. We provide a detailed simulation study which illustrates this below.

In summary, `haploSep` is computationally very efficient, with a linear computational complexity in the number of SNP locations $N$ (up to, potentially, the initial clustering step). In the following we illustrate these computational aspects with simulation examples, which were all performed on a standard laptop with Intel Core i7 processor.

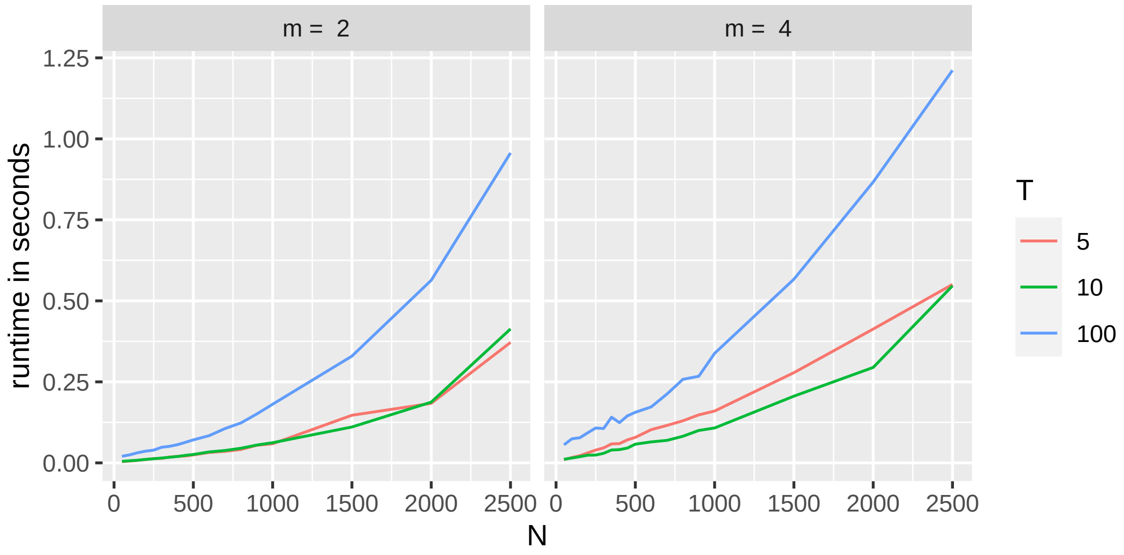


Figure S1: Runtime analysis for `haploSep`. Runtime (y-axis) against number of SNPs $N$, for different values of number of generations $T$ (see legend for color code), and number of reconstructed haplotypes $m$ (left: $m = 2$ and right: $m = 4$). Results are averaged over 100 Monte Carlo runs. See text for details of simulation setup. As can be seen, even for a large number of generations $M$ (e.g., $M = 100$) and a large number of variants (e.g., $N = 2500$), `haploSep` has a runtime of only a few seconds.

For different values of $N, T$, and $m$ we evaluated the run time of `haploSep`. To this end, we randomly generated an $N \times m$ binary matrix as a haplotype structure and then applied the function `haploSimulate` from our R package `haploSep` to simulate an allele frequency matrix $Y \in [0,1]^{N\times T}$ with effective population size equal to 300, at generations $0, 10, 20, \dots, 10\cdot(T-1)$, and with mean sequencing coverage of 80. We took the average over 100 Monte Carlo runs. Results are shown in Fig. S1. As can be seen, even for as many as 100 generations and 2500 variants, `haploSep`'s runtime, e.g., for 4 haplotypes, is just a little bit over a second, which shows that computation time will almost never be problematic for real data applications in a typical sample size regime.

Moreover, we evaluated the number of iterations that `haploSep` performs update stpes of $(\hat{W}, \hat{b})$ and $\hat{S}$, respectively, for different values of stopping thresholds $\delta$. As an example, we considered $N = 500, m = 3, T = 10$, with $Y$ generated in the same way as for the previous simulations. Fig. S2 shows the average number of iterations (y-axis) for $\delta = 10^{-2}, 10^{-3}, \dots, 10^{-10}$ (x-axis) over $1,000$ Monte Carlo runs, with standard deviation shown as error bars. As can be seen, on average `haploSep` performs between 2 and 3 iterations, even when $\delta$ is as small as $10^{-10}$.

To further illustrate robustness with respect to the $\delta$ parameter, we compared the reconstructed $\hat{W}$ and $\hat{S}$ for different values of $\delta$. More precisely, we considered the same simulation setup as before and let $\hat{W}^i, \hat{S}^i$, for $i = 1, 2$ be the reconstruction for $\delta_1 = 10^{-3}$ (which is the default value for our simulations) and $\delta_2 = 10^{-6}$. Fig. S3 shows the mean absolute deviation $|\hat{W}^1_{ij} - \hat{W}^2_{ij}|$ (left) and $|\hat{S}^1_{ni} - \hat{S}^2_{ni}|$ (right) averaged over $i = 1, \dots, 3$, $j = 1, \dots, 10$, $n = 1, \dots, 500$, and $1,000$ Monte Carlo runs with $Y$ as in the previous setup. As can be seen, the difference between $(\hat{W}^1, \hat{S}^1)$ and $(\hat{W}^2, \hat{S}^2)$ is negligible, and hence, we conclude that the choice of $\delta$ is not of major concern.

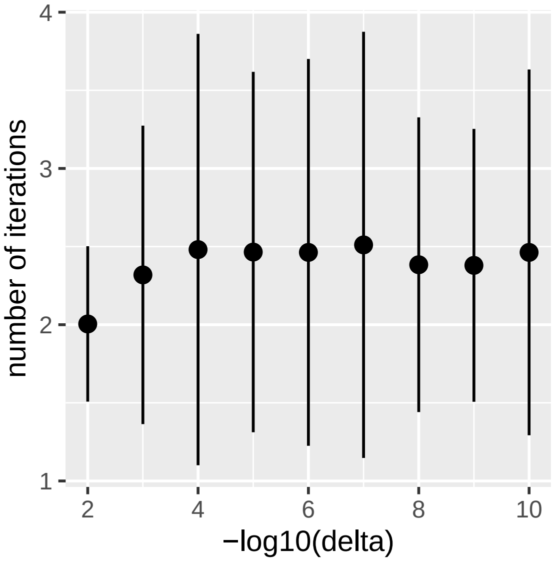


Figure S2: Simulation analysis for the number of iterations that the `haploSep` procedure requires for the Lloyd's-type update setps. Number of iterations (y-axis) for $-\log_{10}(\delta) = 2, 3, \ldots, 10$ (x-axis) averaged over $1,000$ Monte Carlo simulations, with standard deviation shown as error bars. See text for details of simulation setup. As can be seen, `haploSep` typically converges after a few iterations.

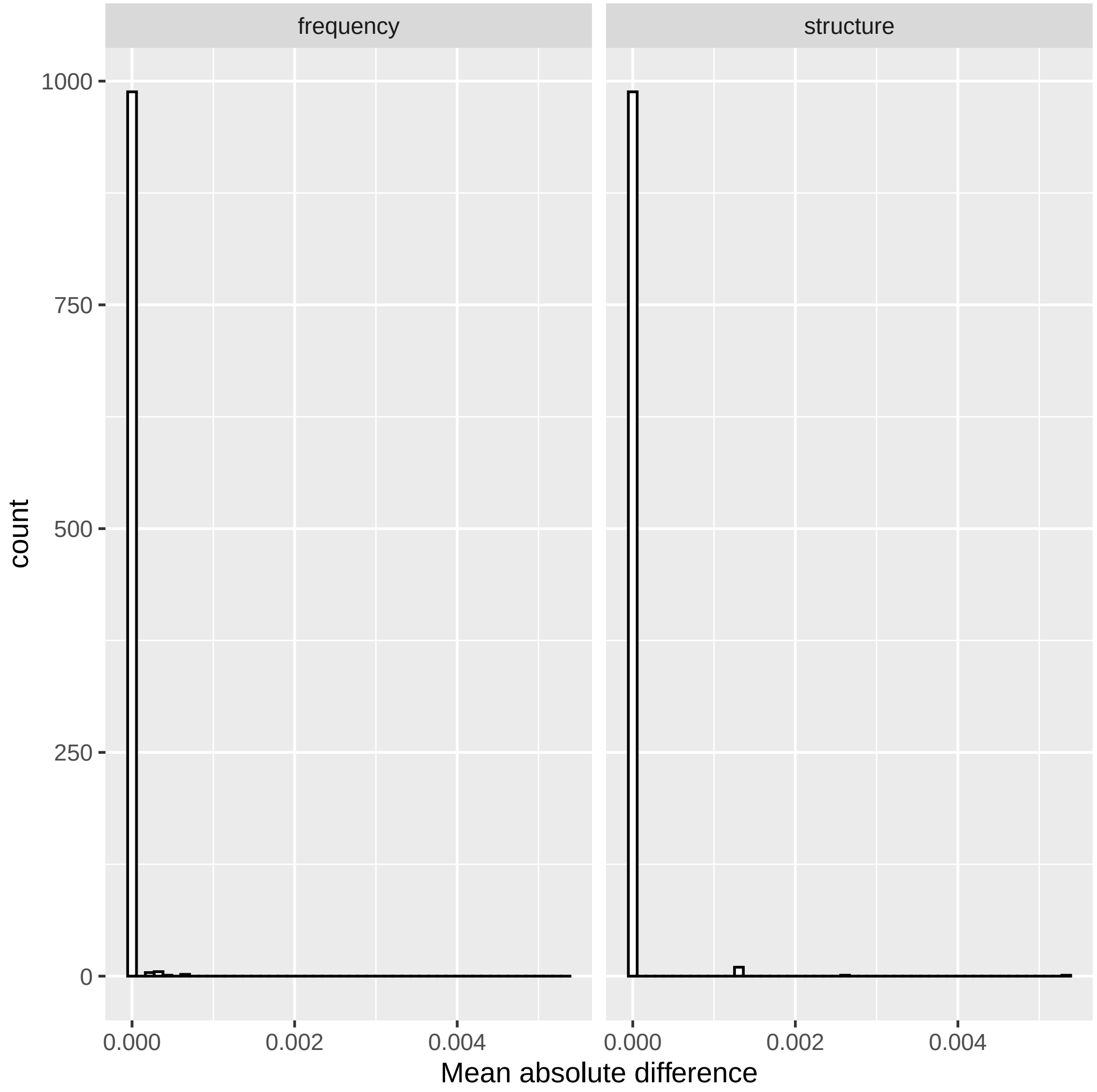


Figure S3: Simulation analysis for the difference in `haploSep`'s reconstruction of $S$ and $W$ for different values of the threshold parameter $\delta$. Histogram of absolute difference of reconstructed frequency matrices, $|\hat{W}^1_{ij} - \hat{W}^2_{ij}|$, (left) and haplotype structure, $|\hat{S}^1_{ni} - \hat{S}^2_{ni}|$, (right) over $i = 1, \ldots, 3$, $j = 1, \ldots, 10$, $n = 1, \ldots, 500$, and $1,000$ Monte Carlo runs with $Y$ as described in the main text. Here, $\hat{W}^i, \hat{S}^i$, for $i = 1, 2$, denotes the reconstruction with threshold value $\delta_1 = 10^{-3}$ (which is our default value) and $\delta_2 = 10^{-6}$, respectively.

#### S2-4 Model selection via SVD

Note that in the noiseless population case ($Y = SW + b$ in (2)) the number of dominant haplotypes $m$ can directly be obtained via the rank of the AF matrix with

$$\operatorname{rank}(SW + \mathbf{1}b^\top) = m + 1. \tag{S3}$$

To see this, note that the $t$th column of $SW + b$ can be written as

$$\sum_{i=1}^{m} S_{\cdot i} W_{it} + b_t (1, \ldots, 1)^\top$$

and thus

$$\operatorname{rank}(SW + \mathbf{1}b^\top) = \dim(\operatorname{span}(S_{\cdot 1}, \ldots, S_{\cdot m}, (1, \ldots, 1)^\top)) = m + 1,$$

where the last equality follows from the identifiability condition (S2). Thus, estimation of $m$ from $Y$ corresponds to estimating the (low) rank of the matrix $SW + b$ from its noisy version $Y$. A more general strategy for the noisy case is to consider the singular values $s_1, \ldots, s_T$ of $Y$ (assuming that $N \geq T$) and then estimate

$$\hat{m} + 1 = \#\{s_i \geq \tau \ : \ i \in [\min(N, T)]\} \tag{S4}$$

for some threshold $\tau$. [Gavish and Donoho, 2014] derived optimal thresholds (in terms of matrix denoising) that are approximately

$$\tau \approx (0.5(T/N)^3 - 0.95(T/N)^2 + 1.82(T/N) + 1.43)\; s_{\text{med}}, \tag{S5}$$

where $s_{\text{med}}$ denotes the median of the singular values $s_1, \ldots, s_T$ of $Y$. In summary, we estimate $\hat{m}$ as in (S4) with $\tau$ as in (S5).

#### S2-5 Accuracy scores

In practice, it may happen that our modeling assumption of a small number of major haplotypes $m \ll T, N$ is violated, e.g., because only few haplotypes are lost over time under some neutral scenario without selection. Alternatively, the selected haplotypes may get lost early on due to random genetic drift. In such a case, a low dimensional haplotype representation will often yield a poor fit to the data $Y$, which we measure using the well known coefficient of determination $R^2 = 1 - \frac{\|Y - \hat{S}\hat{W} - \mathbf{1}\hat{b}^\top\|^2}{\|Y - \bar{Y}\|^2}$. Besides $R^2$, we also report the uncertainty of the proposed estimates via bootstrap confidence scores and bands [Efron, 1979]. Recall that the haplotype structure $S$ is constant over the time points $t \in [T]$. Thus, in order to evaluate uncertainty in the estimate $\hat{S}$, we propose to resample (with replacement) from the empirical distribution on $\{Y_{\cdot 1}, \ldots, Y_{\cdot T}\}$, that is,

$$Y_t^\star \overset{\text{i.i.d.}}{\sim} \frac{1}{T} \sum_{t=1}^{T} 1_{Y_{\cdot t}}, \tag{S6}$$

where $1_y$ denotes the dirac measure on $y$. For each haplotype $i \in [m]$ and SNP location $n \in [N]$ via sampling $Y^\star = (Y_1^\star, \ldots, Y_T^\star)$ from (S6), we compute the variance of $\hat{S}_{ni}(Y^\star)$. As stability score for the $i$th haplotype estimate we report the following score:

$$\text{StabScoreS}_i = 1 - \frac{1}{N} \sum_{n=1}^{N} |\hat{S}_{ni} - \frac{1}{K} \sum_{k=1}^{K} \hat{S}_{kni}| \in [0, 1]. \tag{S7}$$

A stability score of $\text{StabScoreS}_i = 1$ suggestes an unbiased estimate of the $i$th haplotype and stability score of $\text{StabScoreS}_i = 0$ a highly biased estimate, which may occur due to model misspecification (i.e., violation of the major haplotype assumption or the identifiability conditons).

For the haplotype frequencies $W$, we observe that they are invariant for different locations $n \in [N]$. Thus, to evaluate uncertainty for $W$ we resample from

$$Y_n^* \overset{\text{i.i.d.}}{\sim} \frac{1}{N} \sum_{n=1}^{N} 1_{Y_n}. \tag{S8}$$

We report the 0.025 and 0.975 quantiles of $\hat{W}_{it}(Y^*)$ as bootstrap confidence bands and the average width of those confidence bands as stability scores.

In practice, we found the above scores to perform reasonable, but we clearly note that there are many other possibilities to construct quality scores for our setting, such as other bootstrap based scores, or also Bayesian credible scores, or frequentist p-values that are based on explicit modeling assumptions, potentially conditioning on either $\hat{W}$ or $\hat{S}$ to construct conditional confidence statements for the other.

We determine a criterion for accepting scenarios where the reconstruction has enough accuracy overall and consider the structure and frequency specific accuracy scores only for those scenarios. Our criterion is based on the $R^2$ scores and the frequency change of the haplotype reaching highest frequency. More specifically, we require $R^2 > 0.8$ and the frequency change of the haplotype reaching highest frequency $> 0.1$.

### S3 Simulation setup

We evaluate our approach using extensive simulations. In our simulations we considered three experimental designs aiming to reproduce the three data sets we analyze in Section 4, i.e. the experiments explained in [Noble et al., 2019], [Castro et al., 2019] and [Barghi et al., 2019]. They cover three very different organisms used in E&R experiments (*Caenorhabditis elegans*, mice, and *Drosophila simulans*) with various complexities leading to three different starting conditions for the experiments. Indeed, mice populations need to be small because of the maintenance effort involved, whereas this is not the case for *Drosophila simulans* and even less for *Caenorhabditis elegans*. The latter two organisms thus give more freedom to choose the number of different starting haplotypes.

Selection is an important factor in E&R experiments where researchers attempt to understand the genetic architecture of adaptation. In the literature, several E&R experiments have been discussed that involve different stressful conditions. Sources of stress can be high/low-quality food, body size constraints (e.g. only sufficiently small or large organisms are allowed to reproduce), or heat. Our three data sets consider stress conditions on the reproduction regime [Noble et al., 2019], on the body size [Castro et al., 2019] and the temperature regime [Barghi et al., 2019]. Other publications focus on desiccation resistance [Griffin et al., 2017], pathogen resistance [Kraaijeveld and Godfray, 2008], and selection on flying speed [Weber, 1996].

In our simulations, we consider starting populations with the same numbers of haplotypes, and of individuals, as in the real data applications discussed in Section 4. As some of the founder haplotypes from [Barghi et al., 2019] were made available to us by the authors, starting populations were obtained by sampling from these haplotypes. For our basic scenario, we introduce a simple selection regime with selection strength $s = 0.05$ for a beneficial allele present at three different founder haplotypes. The genetic composition of generation $n$ is obtained by multinomial sampling from the previous generation. Sequencing data are generated every tenth generation at 16 different time points ($G_0, G_{10}, \ldots, G_{150}$). From the simulated haplotype data, we compute the true allele frequencies via the regression model $Y = SW$ in Section 2 of the main text as the matrix product of the simulated haplotype structure and frequency. Afterward, we simulate observed allele frequencies using binomial sampling with sample size $n$ equal to the local sequencing coverage, taken from a Poisson(80) distribution. This is to mimic that real allele frequency data in most E&R experiments are noisy because individuals are sequenced as a pool with a given depth (coverage) that changes according to the available resources. With pool sequencing the DNA of all organisms is mixed and sequenced together. An extensive explanation of pool sequencing can be found in [Schlötterer et al., 2014]. A detailed description of this binomial sampling step can be found in [Waples, 1989] and [Jónás et al., 2016].

Beyond our basic scenario, we also investigate several alternative scenarios, and consider how design parameters of E&R experiments affect the quality of our haplotype reconstruction. Parameter values not mentioned in our results have been chosen as in our basic scenario.

### S4 Simulation results

Complementing Section 3.1, we provide results for our three simple selection scenarios on the comparison between the reconstructed and the true haplotype structure in Fig. S4.

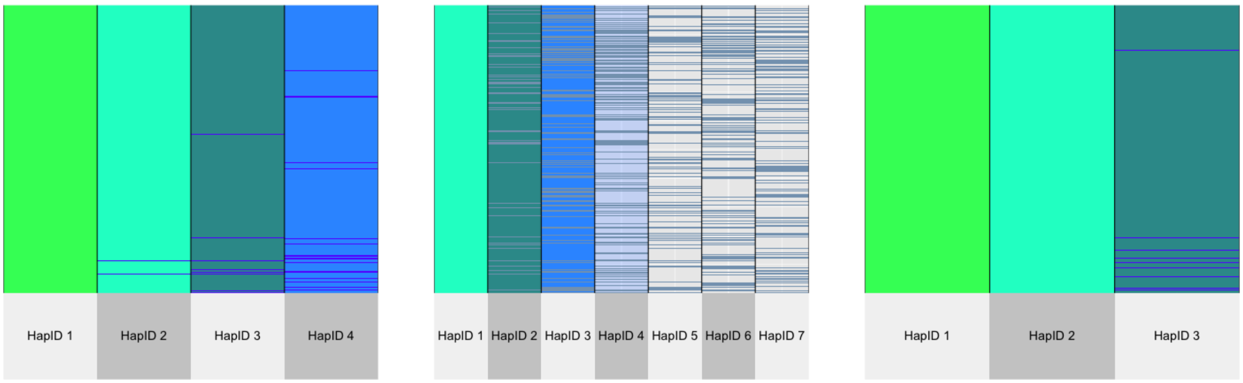


S4a *Longshank mice exp.* S4b *C. elegans* S4c *D. simulans*

Figure S4: Result of one simulation run from the simple selection scenario with the experimental design from the Longshank mice experiment (a), *C. elegans* (b), and *Drosophila simulans* (c). This figure shows inconsistencies between true and reconstructed haplotype structure. Blue line indicates mismatches.

Most of the mismatches that we observe in Fig. S4 are in the low-frequency haplotypes. In order to reconstruct haplotypes correctly, they need to be present in the population at an appreciable frequency for several generations. In particular our approach usually cannot accurately reconstruct the structure of haplotypes reaching zero frequency in the earlier part of the experiment. Even so, those haplotypes are not of interest for most analyses trying to understand the architecture of adaptation because they do not provide any contribution to it. Since the number of true haplotypes can be much larger than the number of haplotypes we reconstruct, we match the (true) haplotype having the closest possible structure to the given reconstructed one to compute the error for our estimated haplotypes. As for the figures in the main text, we filter again using our criteria on $R^2$ and the frequency change of the most abundant haplotype as explained in Section S2-5. See Section S7 for the remaining simulation runs. Based on 100 simulation runs, Fig. S5 shows very low error for both frequency and structure of the selected haplotype(s). However, looking at the different time points, the error is higher for initial generations, whereas it drops for later stages of evolution (see Fig. S5b). The differences between earlier and later time points can be pronounced depending on the experimental design. Indeed when selection occurs, our method provides better estimates for later time points than for earlier ones, if the number of reconstructed haplotypes is much smaller than the number of haplotypes in the starting population. Similar conclusions can be drawn also for the results about the experimental design based on [Noble et al., 2019], shown in Fig. S6.

Starting from these three simple selection scenarios, we did simulations for different values of important parameters for E&R in order to assess how they affect our haplotype reconstruction. We focus on the selection coefficient, the number of haplotypes in the founder population, the number of haplotypes carrying the beneficial allele, the coverage and the number of time points where the sequencing data are collected. For each simulation run the number of haplotypes being reconstructed is estimated via our model selection step as explained in Section S2-4. All the results discussed in this section are simulated with the parameters introduced in Section S3

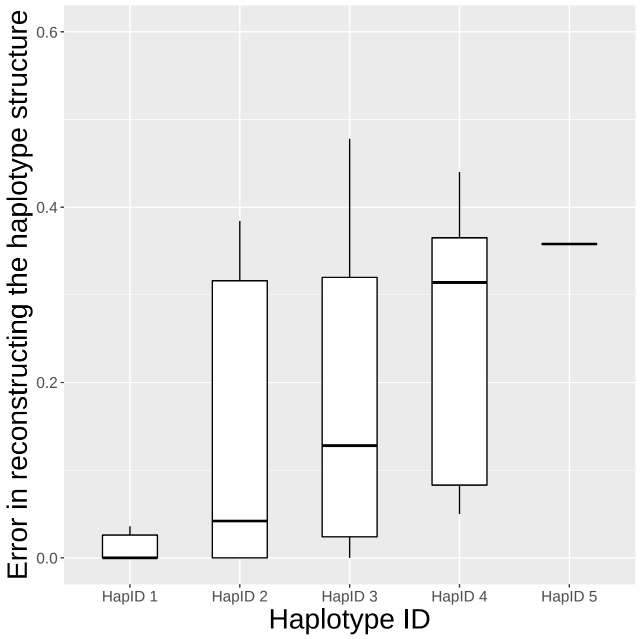


S5a *Structure*

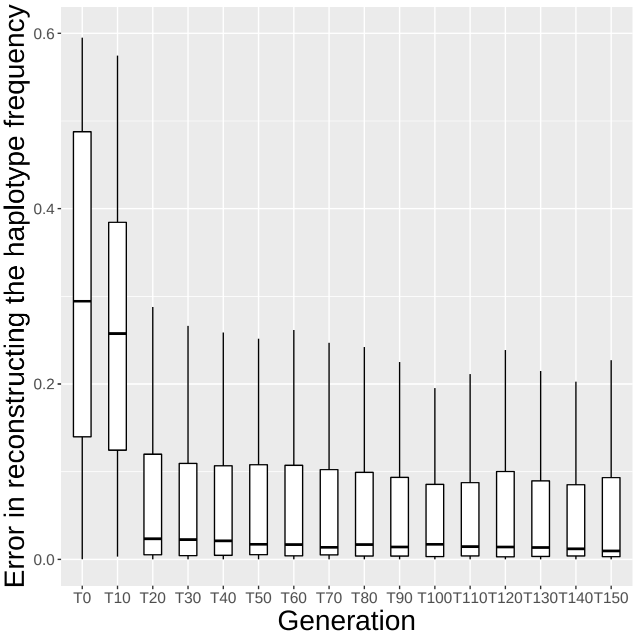


S5b *Frequency*

Figure S5: Haplotype reconstruction error for our basic selection scenario with *Drosophila simulans* based on 100 simulation runs. (a) Proportion of wrongly classified SNPs for each reconstructed haplotype. The haplotypes are displayed in decreasing order according to their cumulative frequency over time. (b) Absolute difference between the true and estimated haplotype frequencies for each time point at which sequencing information is available.

(s = 0.05, 150 generations of E&R where allele frequencies are available every 10 generations, one locus carries the beneficial allele in three individual haplotypes, genotypes from the founder population used in [Barghi et al., 2019]). Fig. S7 shows the accuracy depending on the selection pressure. As we expect, the error decreases when the selection pressure increases. We can observe that the effect is very pronounced for the experimental designs with large population size. This is because the reconstruction results become more and more accurate as the changes in haplotype frequency throughout time increase. When the populations size is small (e.g. in experiments using bigger organisms like mice), these haplotype frequency changes can occur under neutrality as well.

Our method requires information from multiple sources, which for E&R experiments correspond to sequenced time points. The number of time points at which the sequencing data are available mainly depends on the time and costs allocated to the experiment. As it is shown in the lower panel of Fig. S8 (and with a less pronounced effect in the upper panel), four time points do not contain enough information for any experimental design to obtain satisfactory results. However when the number of time points increases the error drops and this is consistent for all three experimental designs as well. It is also important to notice that the number of haplotypes we can reconstruct is smaller or equal to the number of available time points. This can also influence the power of our method under certain experimental designs where a high number of haplotypes is needed to capture the true dynamic of the haplotype frequencies in the given experiment.

In Fig. S9 we consider different numbers of haplotypes sharing the same beneficial allele. The more haplotypes share the same selective advantage, the less accurate the reconstruction becomes, unless the experiment is run for enough time to resolve the competition. If the competition is resolved and one or few haplotype(s) prevail, the reconstruction can reach high accuracy, however.

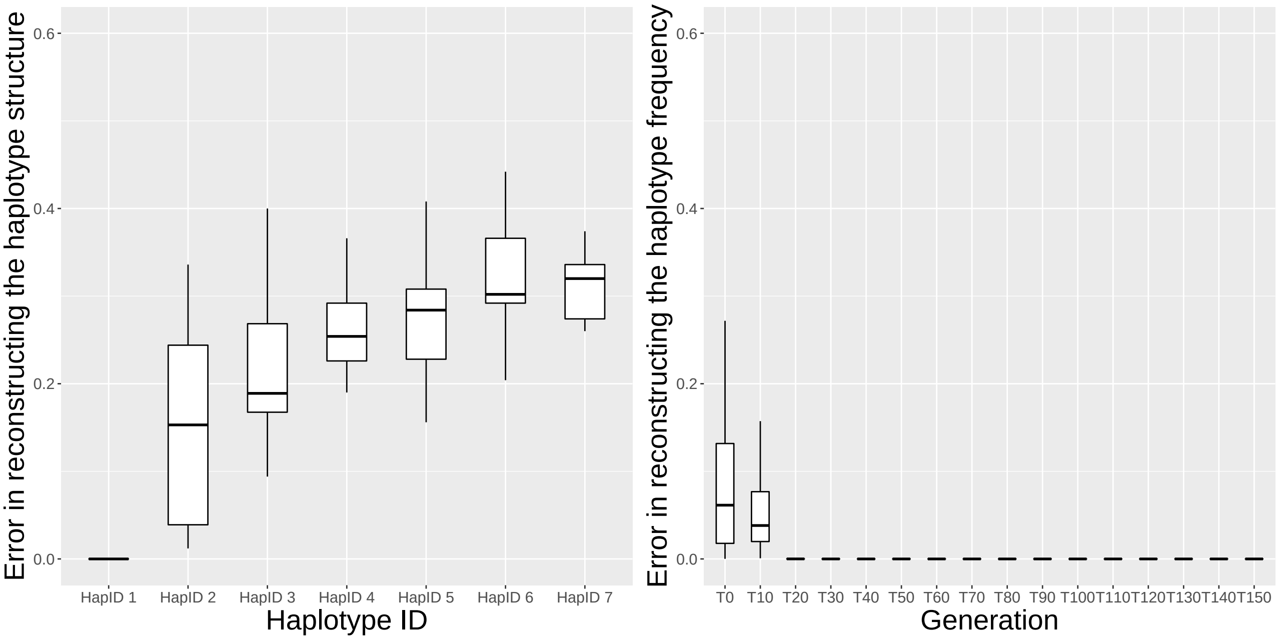


S6a *Structure* S6b *Frequency*

Figure S6: Haplotype reconstruction error for our basic selection scenario with *C. elegans* based on 100 simulation runs. (a) Proportion of wrongly classified SNPs for each reconstructed haplotype. The haplotypes are displayed in decreasing order according to the cumulative frequency over time. (b) Absolute difference between the true and estimated haplotype frequencies for each time point at which sequencing information is available.

When looking at Fig. S10 we can see that a coverage of 5 is too low for accurate pooled allele frequency estimates. Thus our method cannot provide good estimates. When the coverage increases above $\lambda = 20$, not much accuracy is gained anymore. For our considered designs, more time points will be more beneficial than more reads in terms of accuracy. Compare for example, the results from our three experimental designs with fewer time points (e.g. 4) and high coverage ($\lambda = 80$) from Fig. S8 against those with more time point (16) and low coverage (e.g. $\lambda = 20$) from Fig. S10.
The last parameter we considered is the number of different haplotypes in the founder population (Fig. S11). Our simulations do not show a clear trend here. An intermediate number of haplotypes relative to the population size often seems to lead to the highest accuracy, this may be since in this case some - but not all- of the beneficial haplotypes tend to get lost by drift.

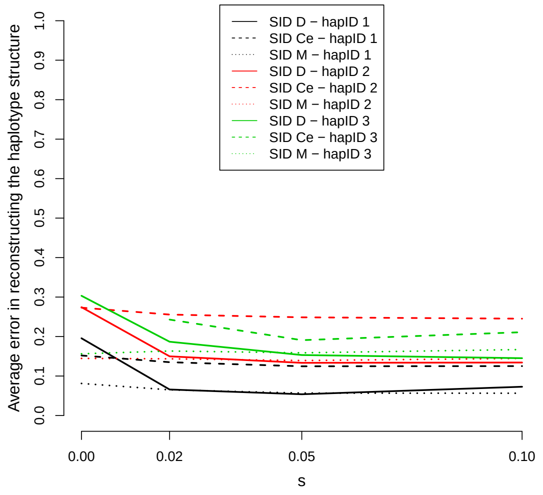


S7a *Structure*

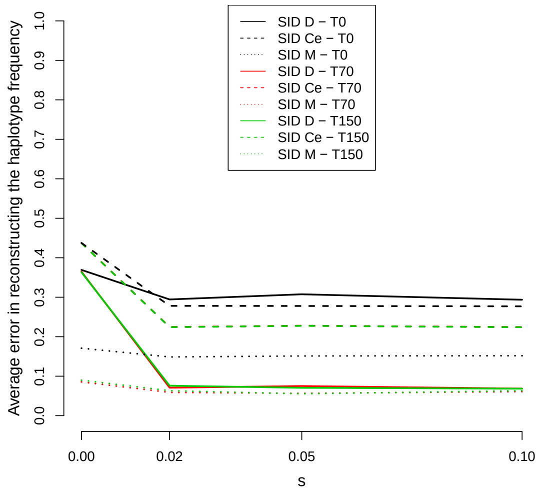


S7b *Frequency*

Figure S7: Dependence of the quality of our reconstruction approach on the selection coefficient. Simulation setup: $s \in 0, 0.02, 0.05, 0.1$ and all the other parameters as in Section S3. Results for *D. simulans* (solid lines), *C. elegans* (dashed lines), and the mice experiment (dotted lines) are shown. (a) Error in reconstructing the haplotype structure versus different values of the selection coefficient. For each experimental design, results for the three most frequent haplotypes are shown: hapID 1 (black lines), hapID 2 (red lines), and hapID 3 (green lines). (b) Error in reconstructing the haplotype frequencies versus different values of the selection coefficient. For each experimental design, results for time points T0 (black lines), T70 (red lines), and T150 (green lines) are shown.

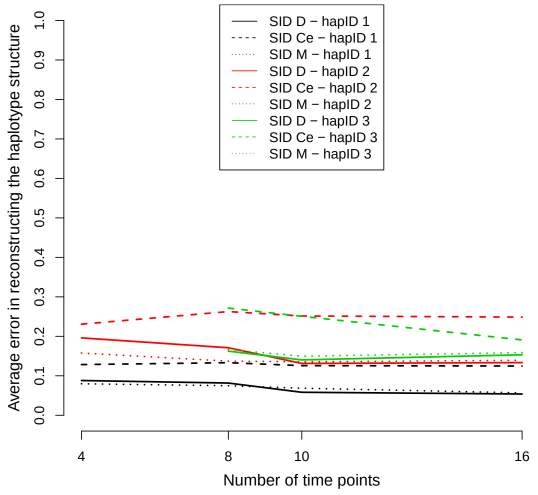


S8a *Structure*

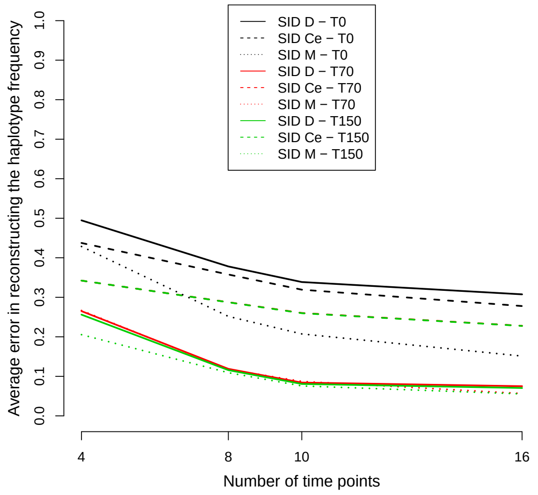


S8b *Frequency*

Figure S8: Dependence of the quality of our reconstruction approach on the number of sequenced time points. Results for *D. simulans* (solid lines), *C. elegans* (dashed lines), and the mice experiment (dotted lines) are shown. (a) Error in reconstructing the haplotype structure versus different numbers of sequenced time points. For each experimental design, results for the three most frequent haplotypes are shown: hapID 1 (black lines), hapID 2 (red lines), and hapID 3 (green lines). (b) Error in reconstructing the haplotype frequencies versus different numbers of sequenced time points. For each experimental design, results for time points T0 (black lines), T70 (red lines), and T150 (green lines) are shown.

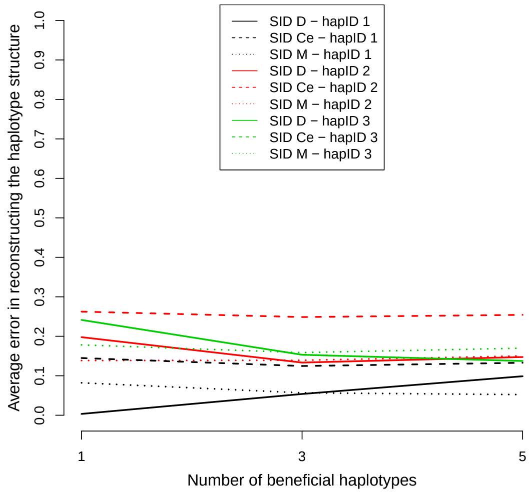


S9a *Structure*

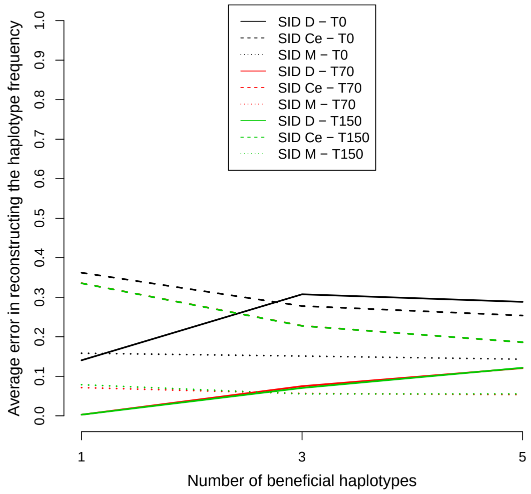


S9b *Frequency*

Figure S9: Dependence of the quality of our reconstruction approach on the number of haplotypes sharing the beneficial allele. Simulation setup: Number of haplotypes sharing the beneficial allele $\in 1, 3, 5$ and all the other parameters as in Section S3. Results for *D. simulans* (solid lines), *C. elegans* (dashed lines), and the mice experiment (dotted lines) are shown. (a) Error in reconstructing the haplotype structure versus different numbers of haplotypes sharing the beneficial allele. For each experimental design, results for the three most frequent haplotypes are shown: hapID 1 (black lines), hapID 2 (red lines), and hapID 3 (green lines). (b) Error in reconstructing the haplotype frequencies versus different numbers of haplotypes sharing the beneficial allele. For each experimental design, results for time points T0 (black lines), T70 (red lines), and T150 (green lines) are shown.

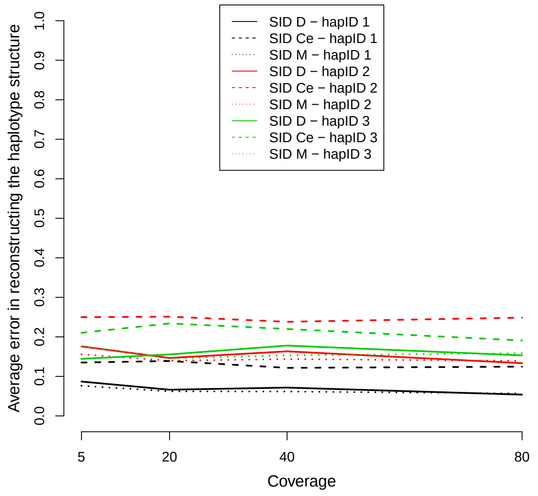


S10a *Structure*

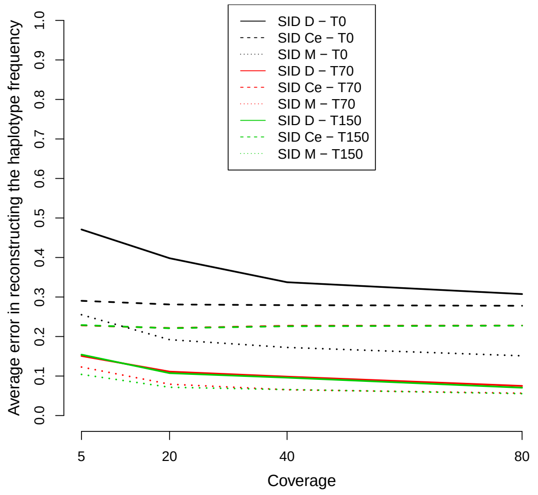


S10b *Frequency*

Figure S10: Dependence of the quality of our reconstruction approach on the mean coverage value $\lambda$. Simulation setup: $\lambda \in 5, 20, 40, 80$ and all the other parameters as in Section S3. Results for *D. simulans* (solid lines), *C. elegans* (dashed lines), and the mice experiment (dotted lines) are shown. (a) Error in reconstructing the haplotype structure versus different values of $\lambda$. For each experimental design, results for the three most frequent haplotypes are shown: hapID 1 (black lines), hapID 2 (red lines), and hapID 3 (green lines). (b) Error in reconstructing the haplotype frequencies versus different values of $\lambda$. For each experimental design, results for time points T0 (black lines), T70 (red lines), and T150 (green lines) are shown.

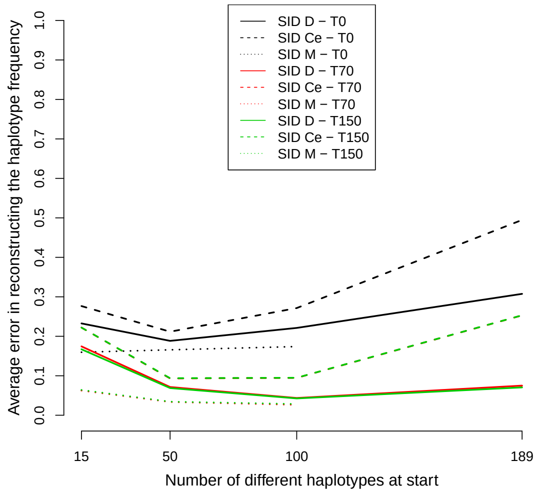


S11a *Structure*

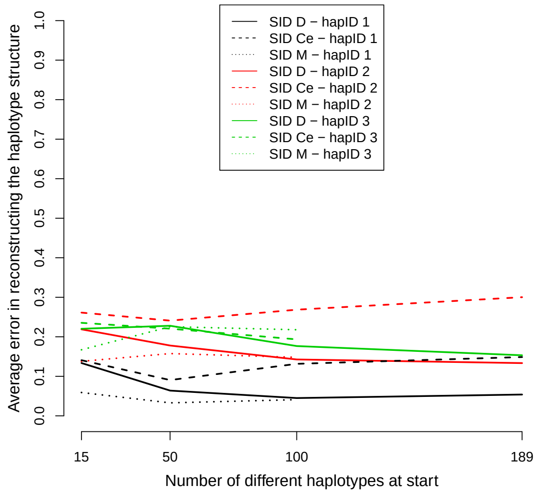


S11b *Frequency*

Figure S11: Dependence of the quality of our reconstruction approach on the number of different haplotypes in the founder population. Simulation setup: Number of different haplotypes in the founder population $\in 15, 50, 100, 189$ and all the other parameters as in Section S3. Results for *D. simulans* (solid lines), *C. elegans* (dashed lines), and the mice experiment (dotted lines) are shown. (a) Error in reconstructing the haplotype structure versus different number of different haplotypes in the starting population. For each experimental design, results for the three most frequent haplotypes are shown: hapID 1 (black lines), hapID 2 (red lines), and hapID 3 (green lines). (b) Error in reconstructing the haplotype frequencies versus different number of different haplotypes in the starting population. For each experimental design, results for time points T0 (black lines), T70 (red lines), and T150 (green lines) are shown.

### S5 Accuracy measures

When applying our method to real data the true haplotypes are unknown and the error cannot be assessed. For this reason, we provide measures of accuracy for the full reconstruction (namely $R^2$) for the haplotype structures and for the haplotype frequencies (see Section S2-5 for a more detailed explanation on how the scores are computed). To see how well these accuracy measures coincide with the actual amount of error, we provide simulation results for our three simple selection scenarios. We expect high scores when the error is low and vice-versa.
We plot $R^2$ against the overall error in the reconstruction of the haplotype frequency for our three simple selection scenarios in Fig. S12. This figure shows that for the scenario with small population size the correlation between $R^2$ and error is relatively high (0.799), however for large population sizes either the correlation is low (0.421) or the $R^2$ is underestimating our error in reconstruction (see Fig. S12c). When the correlation is low, the error is only slightly over estimated by $R^2$, whereas in the case of Fig. S12c we have a group of scenarios where the $R^2$ is too liberal. However, if we discard the scenarios where the haplotype frequency change of the most frequent reconstructed haplotype is small ($< 0.1$) then the correlation in Fig. S12b increases up to 0.521 and the scenarios where $R^2$ is underestimating the error in S12c are not included in the analysis anymore. If the frequency change of the dominant haplotype is small, it means that selection is either not present (neutral dynamic in a large population), or its signal cannot be captured by our method. Therefore we recommend to look at the combination of both $R^2$ and frequency change. This was the motivation for our filtering criteria proposed in Section S2-5.
Our structure specific stability score (see equation S7 in section S2-5) is also correlated with the error in the reconstructed haplotype configuration (see Figs. S13a, S14a, and S15a). The high correlation shows that this measure is useful in applications. To test our accuracy measure for the haplotype frequencies, we checked how often each true frequency is contained inside the accuracy interval. The results in Fig. S16 show a high match between our bands and the true haplotypes, especially for late time points. Histograms of band sizes for these three scenarios can be found in figures S13b, S14b, and S15b, and they reveal that the bands are usually quite small (about 50% or more of the observed bandwidth being smaller than 0.05 in the worst scenario). These results demonstrate that these scores are concordant with the actual errors.
We recommend to use the haplotype specific stability intervals and stability scores after ensuring that our overall quality measures ($R^2$ and frequency change of the dominant haplotype) are good enough.

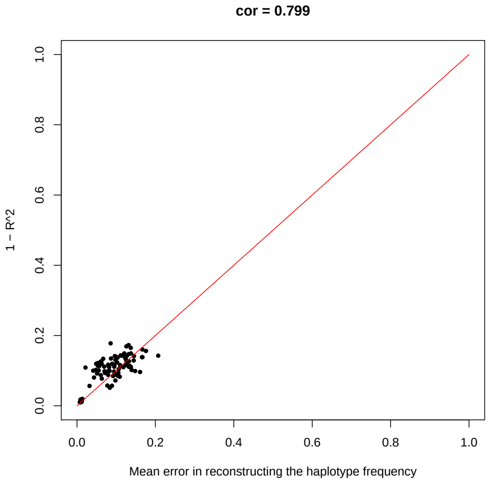


S12a *Longshank mice exp.*

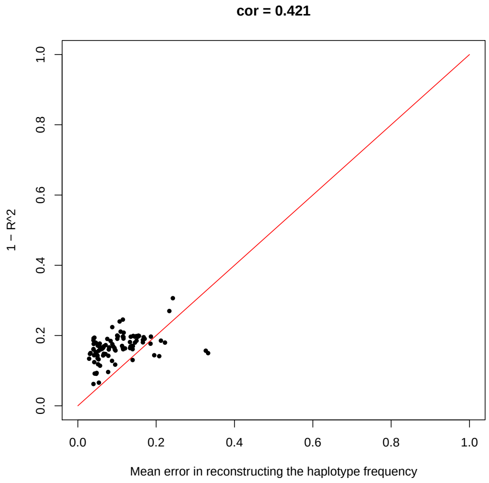


S12b *D. simulans*

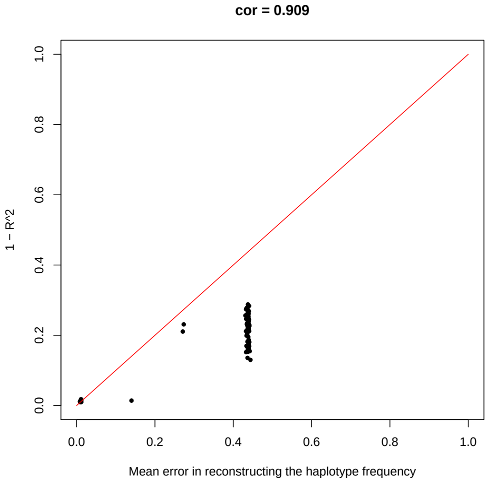


S12c *C. elegans*

Figure S12: Mean error in reconstructing the haplotype frequency versus $1 - R^2$ for (a) the Longshank mice experimental design, (b) the *D. simulans* experimental design, and (c) the *C. elegans* experimental design

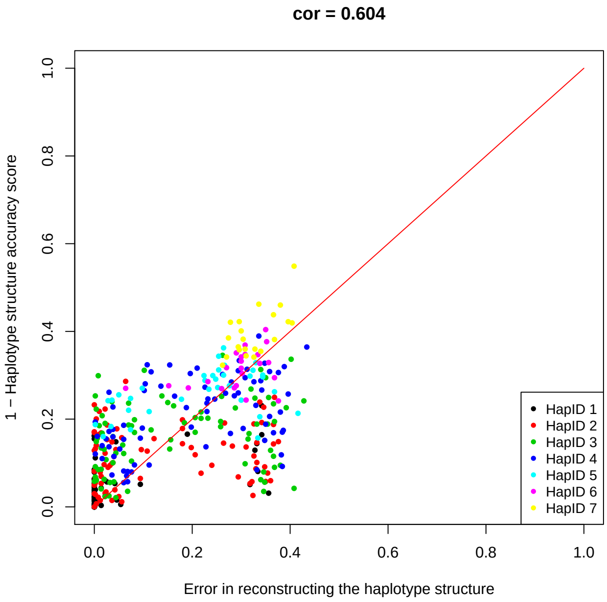


S13a *Structure*

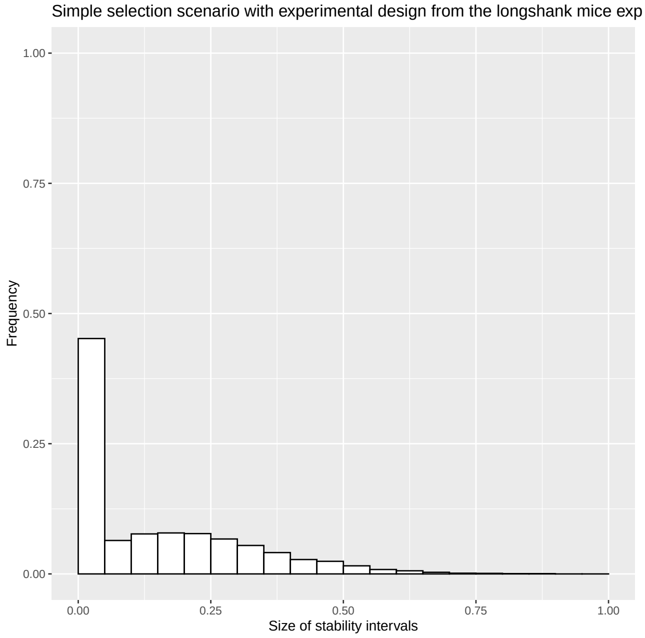


S13b *Frequency*

Figure S13: (a) Proportion of incorrectly estimated alleles when reconstructing the haplotype structure versus the corresponding accuracy scores for the Longshank mice experimental design. (b) Size of the accuracy intervals for the reconstructed haplotype frequency for the Longshank mice experimental design.

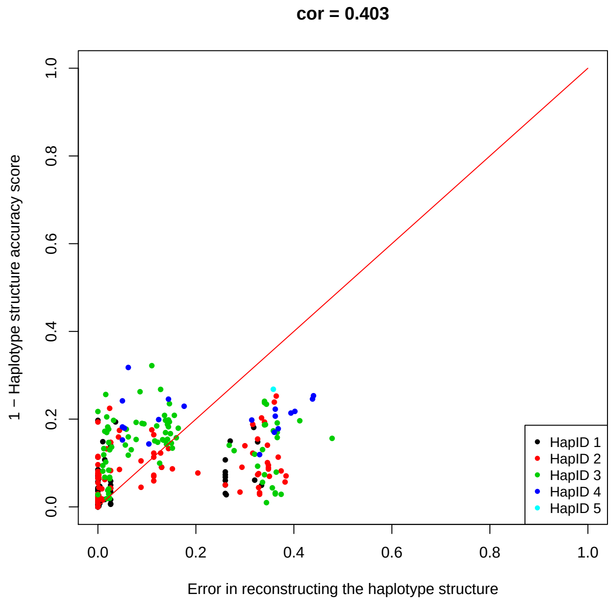


S14a *Structure*

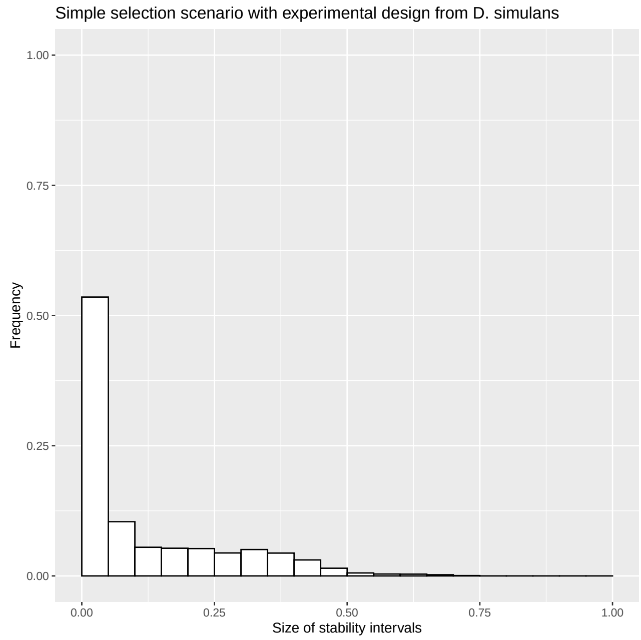


S14b *Frequency*

Figure S14: (a) Proportion of incorrectly estimated alleles when reconstructing the haplotype structure versus the corresponding accuracy scores for the *D. simulans* experimental design. (b) Size of the accuracy intervals for the reconstructed haplotype frequency for the *D. simulans* experimental design.

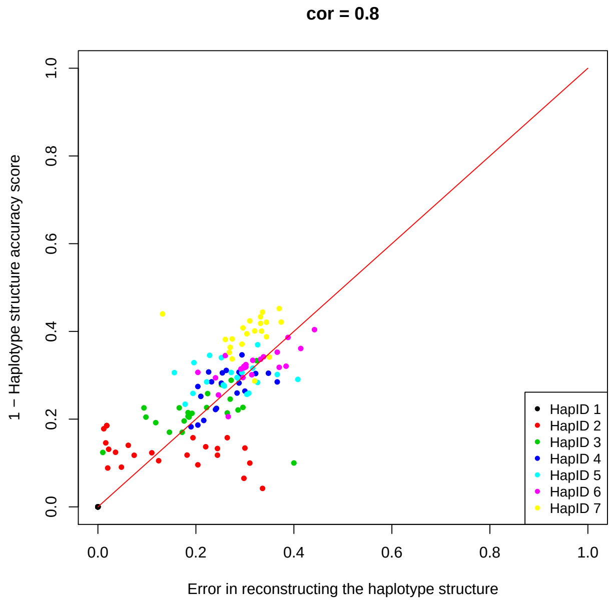


S15a *Structure*

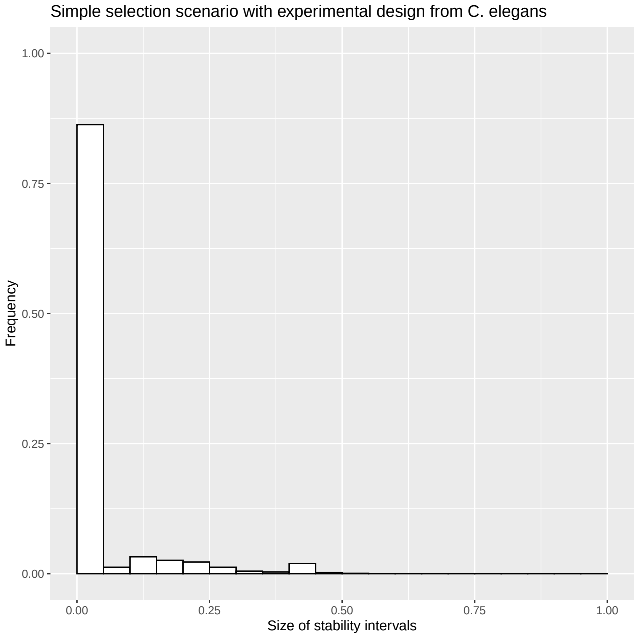


S15b *Frequency*

Figure S15: (a) Proportion of incorrectly estimated alleles when reconstructing the haplotype structure versus the corresponding accuracy scores for the *C. elegans* experimental design. (b) Size of the accuracy intervals for the reconstructed haplotype frequency for the *C. elegans* experimental design.

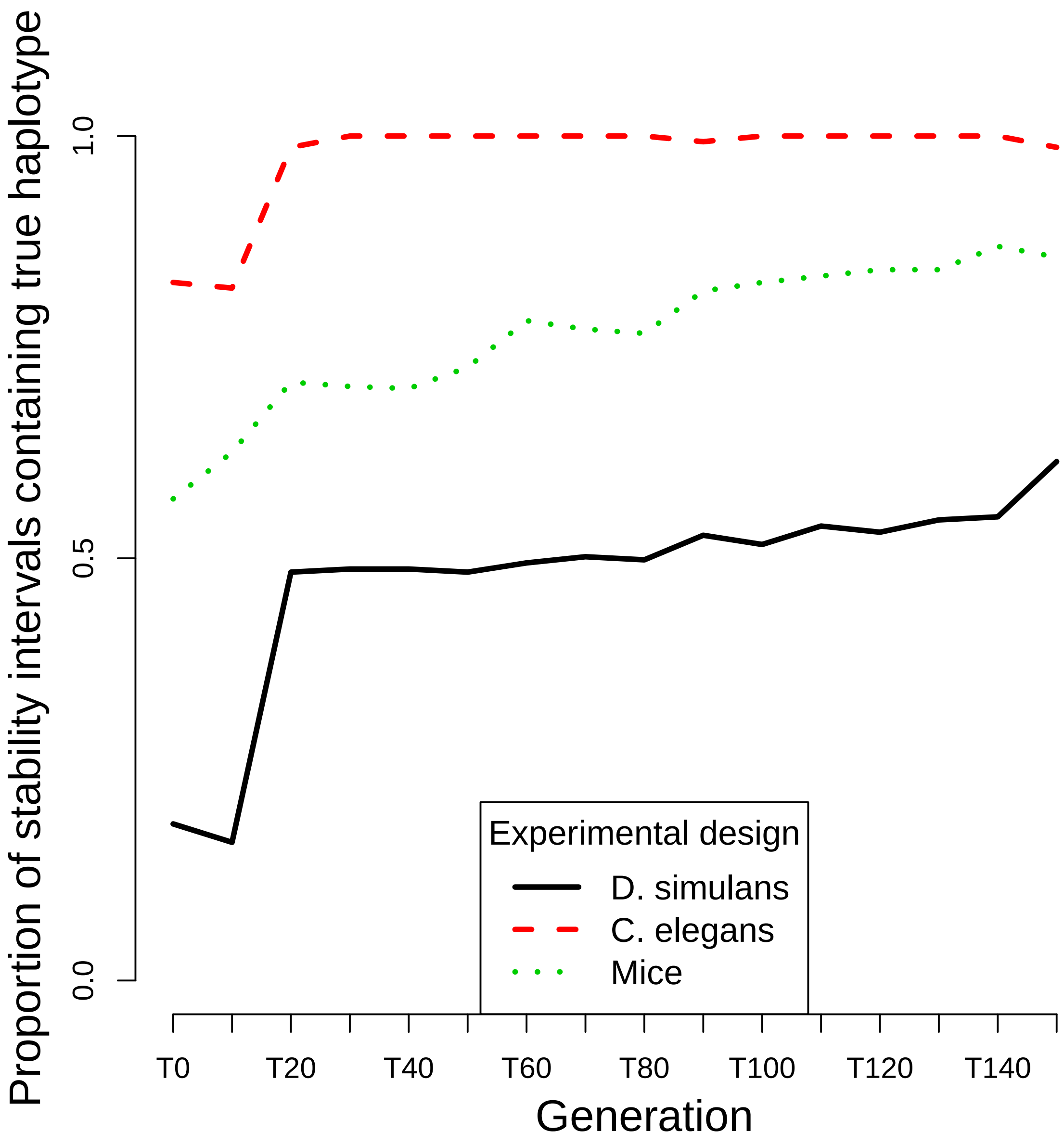


Figure S16: Proportion of stability intervals containing the true haplotype for our three simple selection scenarios.

### S6 Improved allele frequency estimates: additional results

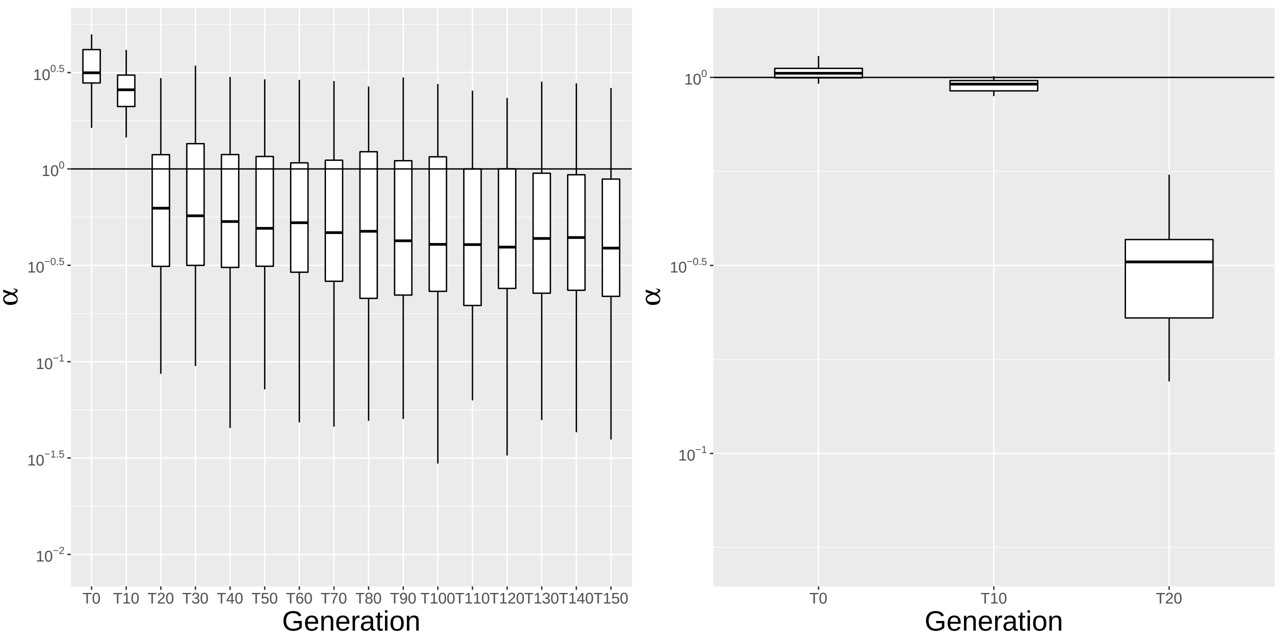


S17a *D. simulans* S17b *C. elegans*

Figure S17: Error ratio ($\alpha$) between haplotype based allele frequency estimates (numerator) and the pool sequencing estimates (denominator) plotted on a log-scale. Results from 100 simulation runs based on the experimental designs in [Barghi et al., 2019] and [Noble et al., 2019].

Late time points for the *C. elegans* example are not shown as both errors in reconstructing the allele frequency data are negligible and thus the ratio cannot be computed. Further information on the later time point can be found in Fig. S20c where all scenarios are included.

### S7 Analysis of outliers

Here we consider all the simulation results for the three simple selection scenarios without filtering using $R^2$ and the frequency change of the haplotype with highest frequency. Figs. S18, S19, and S20 show the quantiles of the errors in reconstructing the haplotype frequency and structure and for $\alpha$. The proportion of scenarios leading to outliers in the error measurements is $15\%, 19\%$, and $78\%$ for the simulations based on the *Drosophila simulans*, Longshank mice, and *C. elegans* experimental design respectively. For *C. elegans* the proportion of outlier simulation runs is considerably higher than for the other two scenarios. Indeed, the population size in the *C. elegans* experiment is much larger than for the other organisms. When the dynamic is neutral in such a large population, there is a large number of haplotypes at very low frequency. These haplotypes are often aggregated within a few estimates at intermediate (and constant) frequency.

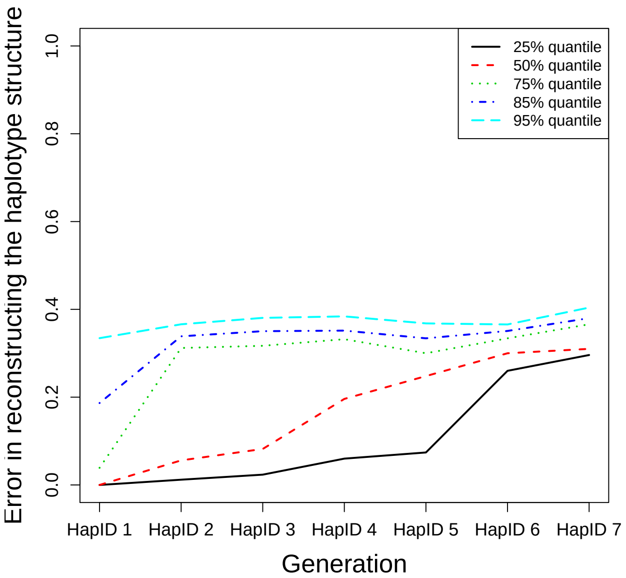


S18a *Longshank mice exp.*

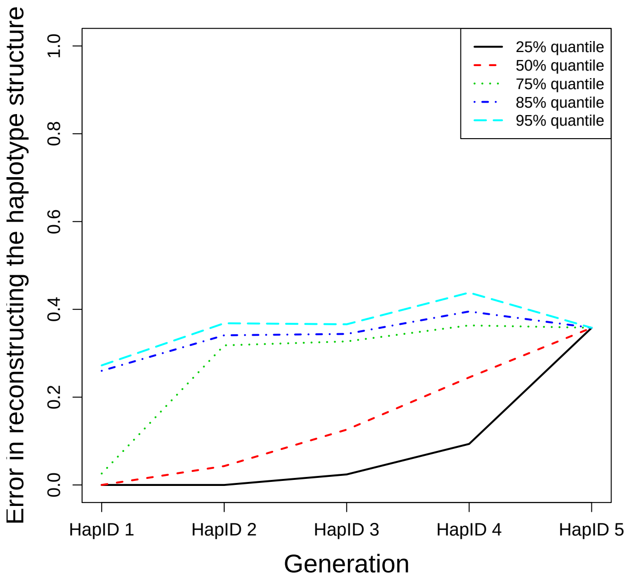


S18b *D. simulans*

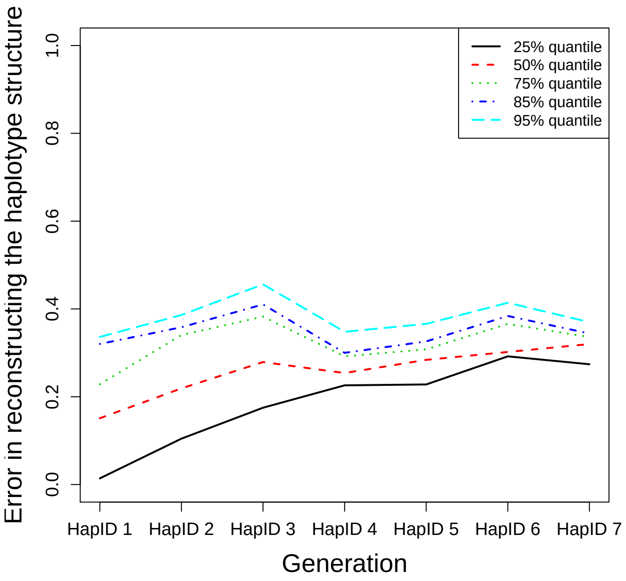


S18c *C. elegans*

Figure S18: Quantiles of the error in reconstructing the haplotype structure for (a) the Longshank mice experimental design, (b) the *D. simulans* experimental design, and (c) the *C. elegans* experimental design.

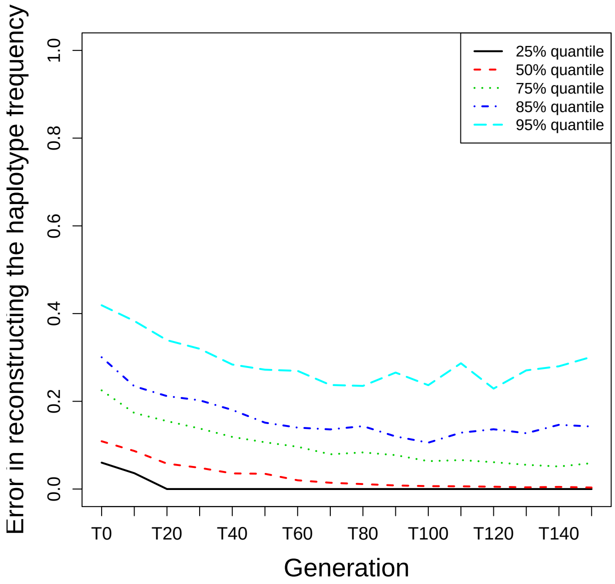


S19a *Longshank mice exp.*

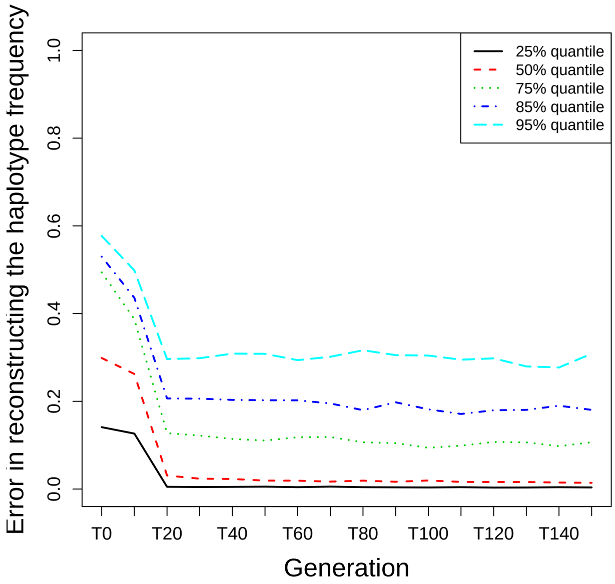


S19b *D. simulans*

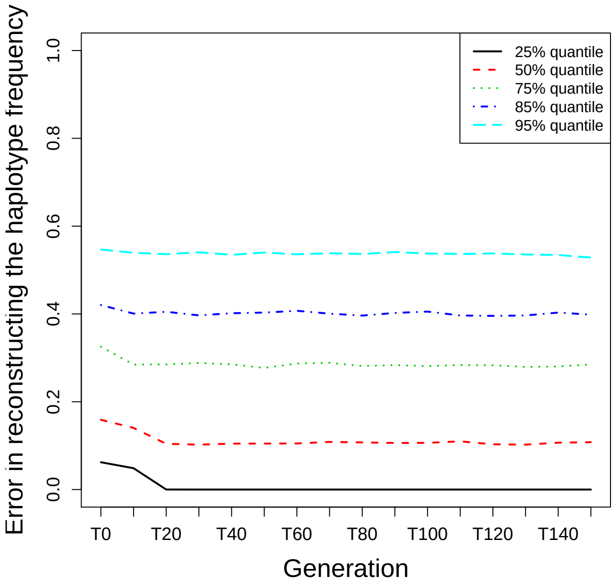


S19c *C. elegans*

Figure S19: Quantiles of the error in reconstructing the haplotype frequency for (a) the Longshank mice experimental design, (b) the *D. simulans* experimental design, and (c) the *C. elegans* experimental design.

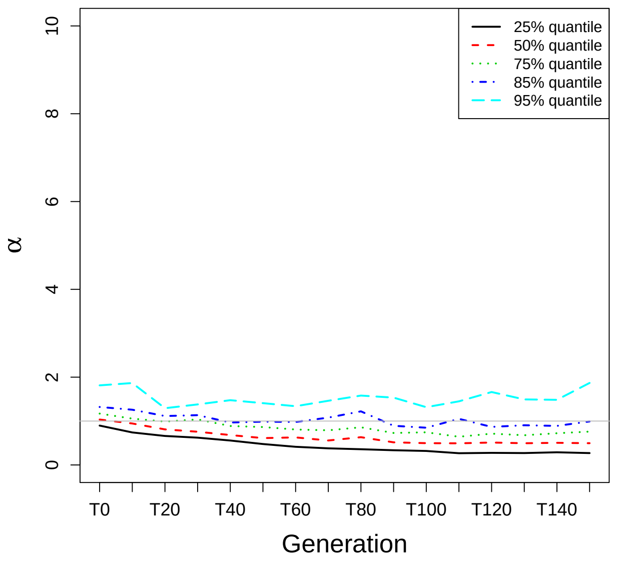


S20a *Longshank mice exp.*

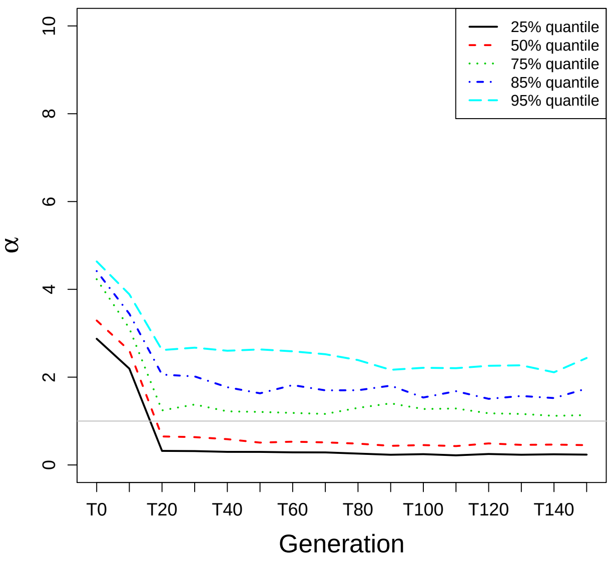


S20b *D. simulans*

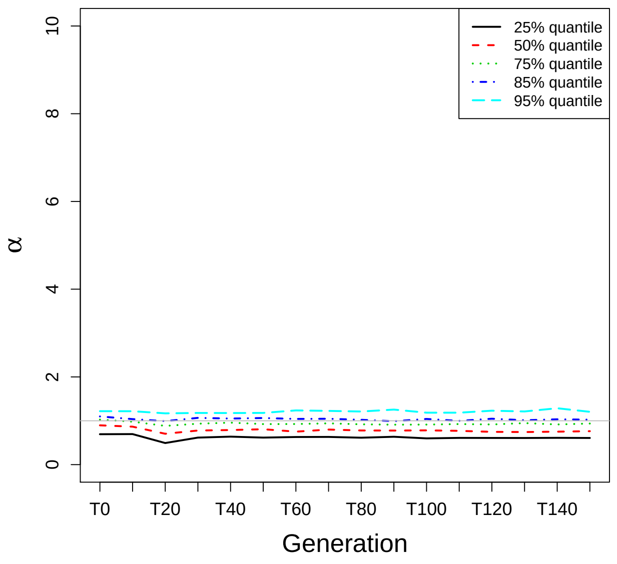


S20c *C. elegans*

Figure S20: Quantiles of the ratio between the error in estimating the allele frequencies from the reconstructing haplotypes versus pool sequencing ($\alpha$) for (a) the Longshank mice experimental design, (b) the *D. simulans* experimental design, and (c) the *C. elegans* experimental design.

### S8 Recombination

We simulated data involving recombination with MimicrEE2 [Vlachos and Kofler, 2018]. For this purpose, we considered a population of 500 individuals with 10 founder haplotypes and a range of recombination rates between 0 and 20 cM/Mb. We used a sample from the *Drosophila melanogaster* genetic reference panel [MacKay et al., 2012] to build our starting population. We ran our simulations for 150 generations assuming that there are three SNPs each having a beneficial allele under selection of strength $s = 0.1$. High recombination rates increase the probability of recombination events putting beneficial SNPs on a new more beneficial haplotype that could rise then considerably in frequency. We store data every tenth generation at time points $G_0, G_{10}, \ldots, G_{150}$. After obtaining the simulated populations, we add sequencing noise to our allele frequency data via binomial sampling under a Poisson coverage with mean 80. The recombination rate has been assumed to be homogeneous throughout the whole region. As in Section 3.1, we performed 100 simulation runs per recombination rate. The results were filtered and those kept for further analysis, where $R^2 > 0.8$ and a frequency change of more than 0.1 was observed for the most frequent haplotype. Our recombination rate in cM/Mb is converted by MimicrEE2 to a lambda-value of a Poisson distribution using Haldane's map function.

The figures below summarize our simulation results and are discussed in Section 3.3.

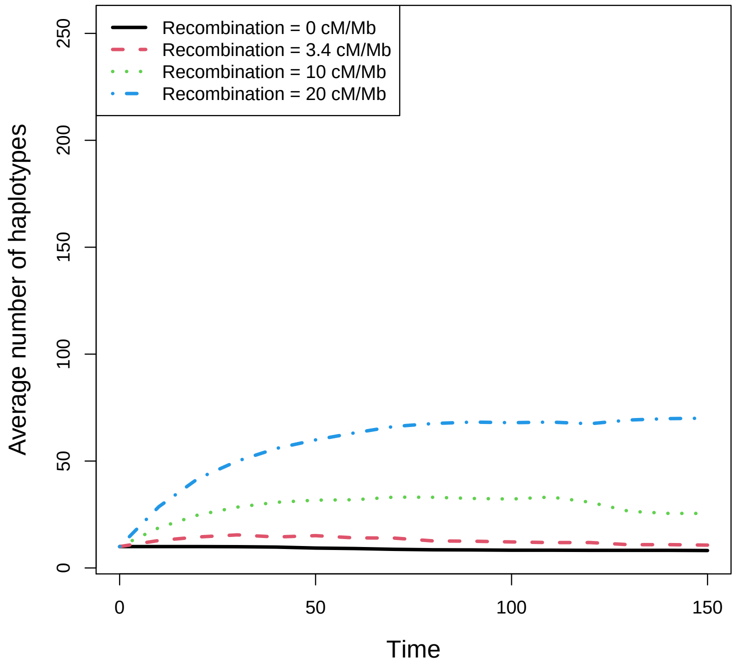


Figure S21: Mean number of haplotypes present at the time points $\{0, 10, 20, \ldots, 150\}$. Each color represents a different recombination rate. For details on this simulation experiment, see the text in this Section.

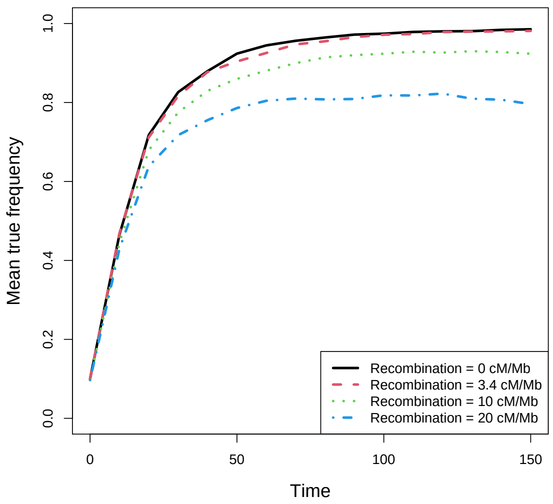


S22a

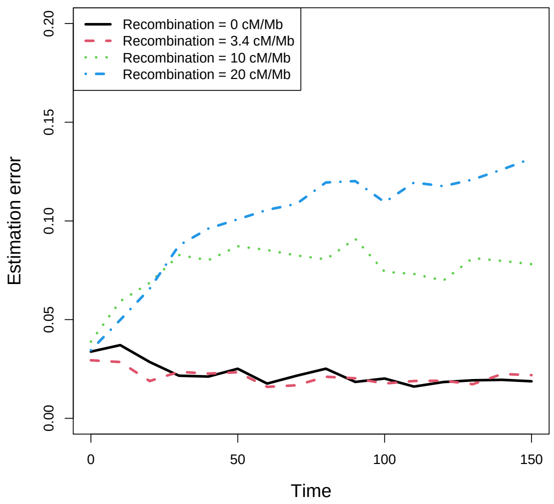


S22b

Figure S22: Frequency of the most abundant haplotype. (a) True frequency of the most frequent reconstructed haplotype: For the most abundant estimated haplotype, the frequencies of the matching true haplotype are averaged across all simulation runs. (b) Estimation error for the most frequent haplotype: Average (across simulation runs) absolute difference between the true and estimated haplotype frequencies for each time point at which sequencing information is available. Different colors and line types indicate the recombination rates $r \in \{0, 3.4, 10, 20\}$.

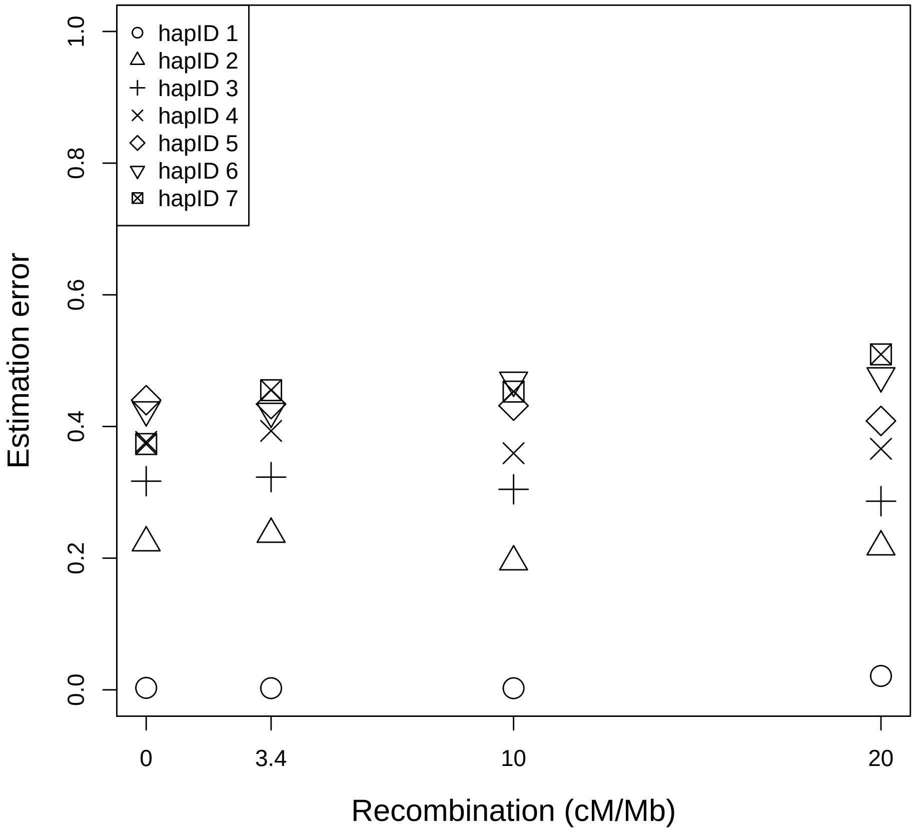


Figure S23: Errors in the estimated haplotype structures for different values of the recombination rate: Proportion of wrongly classified SNPs averaged across simulation runs. Each reconstructed haplotype is matched to the closest true one. The haplotypes are numbered in decreasing order according to their cumulative estimated frequency.

### S9 Validation of our results using read data

We used read data from [Barghi et al., 2019] as a further validation of our reconstructed haplotypes. These data are provided by the authors after the reads were trimmed and mapped to the genome and after duplicates have been removed. These steps, as well as the DNA extraction and library preparation are described in [Barghi et al., 2019]. In order to be consistent with the allele frequency data and thus with the reconstructed haplotypes we only used SNPs analysed in the original paper. Furthermore, as in [Barghi et al., 2019] for a given SNP we kept the information from the reads only when the respective base quality score was higher than 20. As in Section 4, for this analysis we chose a region under selection according to the p-values from the modified chi-squared test in [Spitzer et al., 2020]. Here, we considered the region from 11.239636 to 11.733131 Mb of chromosome 2L in replicate three. All comparisons with the reads are performed at generation 60.

For each read partially overlapping the region of interest we apply the following steps. First, we combined paired end reads to a long sequence with a missing part in the middle because read pairs belong to the same haplotype. Then, we polarize the set of read data for the rising allele, as we did for the allele frequency data.

In order to compare the read data with the reconstructed haplotypes, we considered sliding windows of 1000 SNPs and performed the following analysis on each window. For our first comparison, we selected the most similar read for each reconstructed haplotype and window. Fig. S24 shows the proportion of mismatches between haplotype and corresponding read without considering missing data. From the example we can see that most haplotypes have a good match with the reads, which is a further validation of the fact that the haplotype structure we reconstruct with our method is accurate. However, the number of positions entering this comparison for each read is limited (between 32 and 59). Indeed, there are always many missing values in each read as read length is limited and they might not overlap a region entirely and genomic positions might be filtered out for low base quality scores.

We decided then to examine these results in terms of haplotype frequency as well. Because reads are short and insert sizes generate missing values, we cannot compare the frequencies of the reads with those of the haplotypes directly. At the same time, using single SNPs would not be informative in this situation because we already validated the power of our method in reconstructing allele frequency data (see Section 3.2). Thus, we decided to consider the smallest available linked unit, and we performed our comparison on pairs of subsequent SNPs using the frequencies of the four possible genotypes of each pair.
The results from this comparison are shown in Fig. S25. From these examples we can see that also the frequency of the pairs of SNPs are estimated with low error from our reconstructed haplotypes, which strongly suggests that the reconstructed haplotypes capture the signal from the true haplotypes in the population correctly.

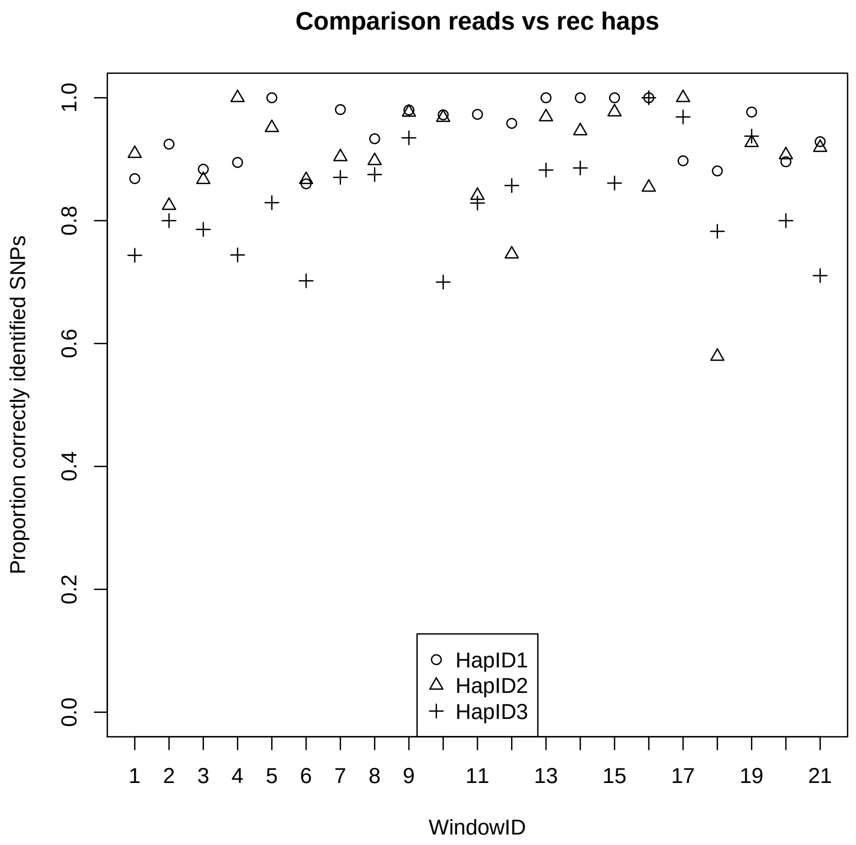


Figure S24: Comparison of the reconstructed haplotype structure with the read data.

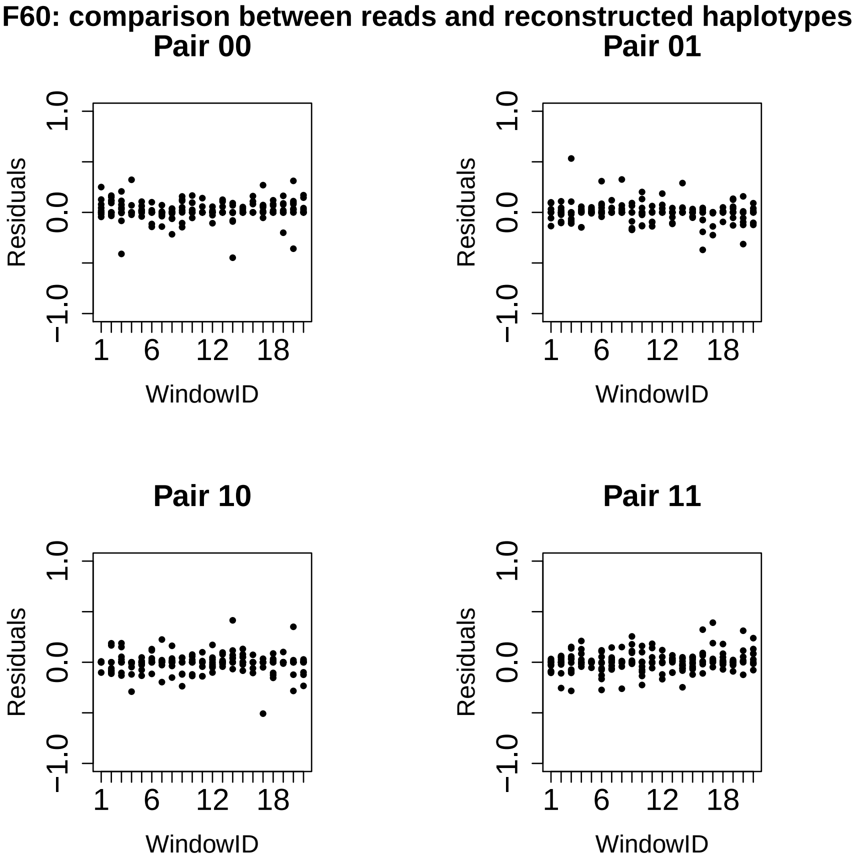


Figure S25: Residuals of the estimated frequency of pairs of SNPs from read data versus the estimated frequency of pairs of SNPs from reconstructed haplotypes.

### S10 Results for the Longshank mice experiment

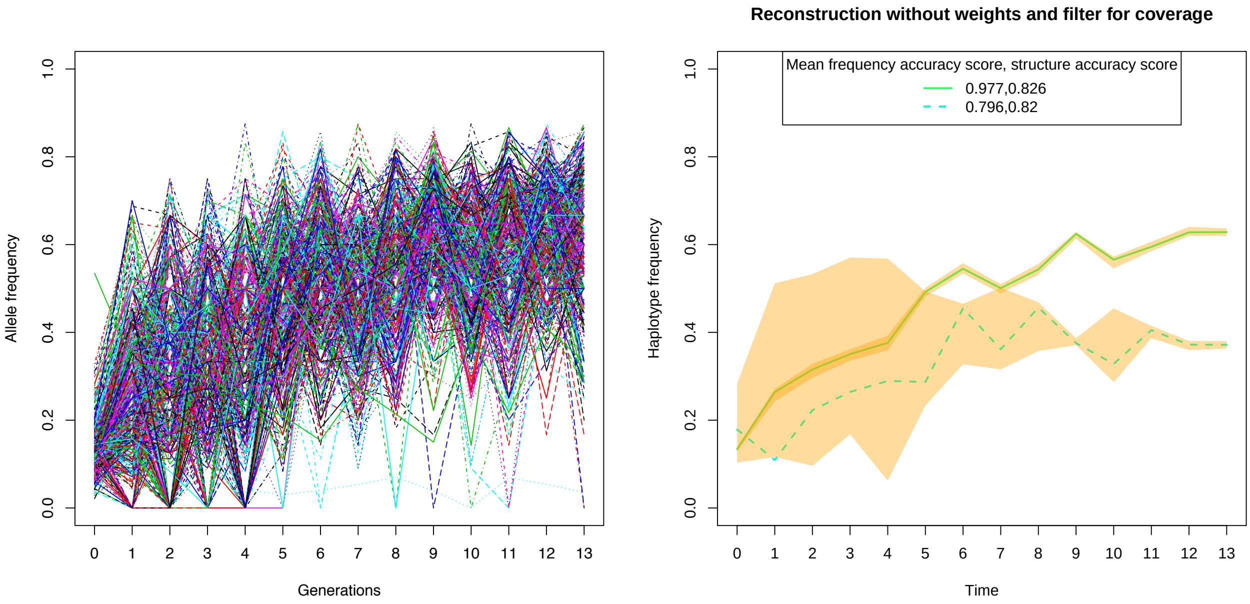


S26a *Alleles* S26b *Haplotypes*

Figure S26: (a) Observed time-series of allele frequencies. (b) Reconstructed haplotype frequencies with accuracy intervals (in yellow) and mean accuracy scores.

### S11 Additional results from the *C. elegans* data set from [Noble et al., 2019]

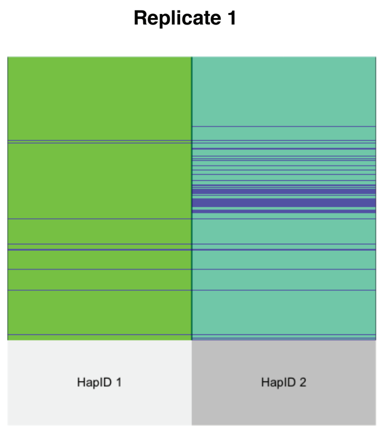


S27a *Structure*

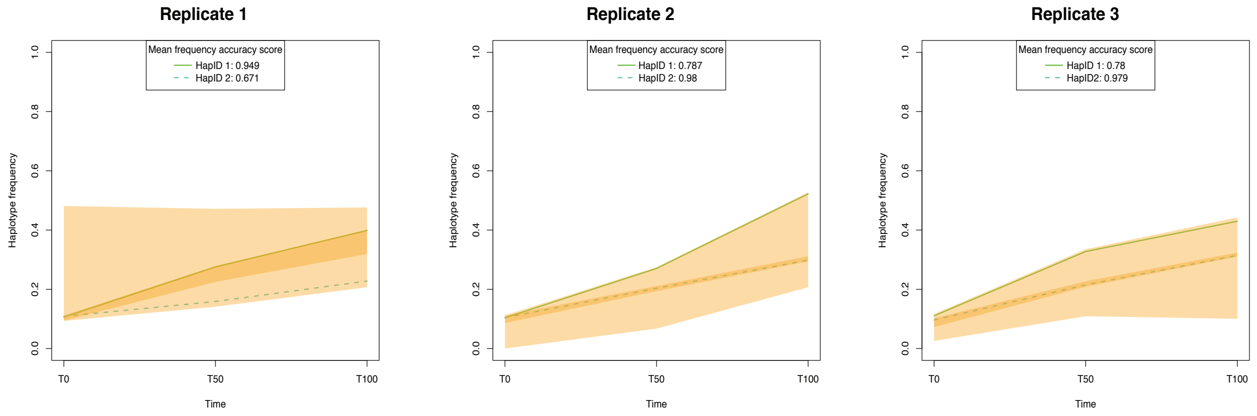


S27b *Frequency*

Figure S27: Haplotype reconstruction for data from [Noble et al., 2019] (a) Match between the haplotype structure reconstructed from the allele frequency data and the sequenced founder haplotypes. Blue lines indicate mismatch positions. (b) Reconstructed haplotype frequencies with accuracy intervals (in yellow) and mean accuracy scores.

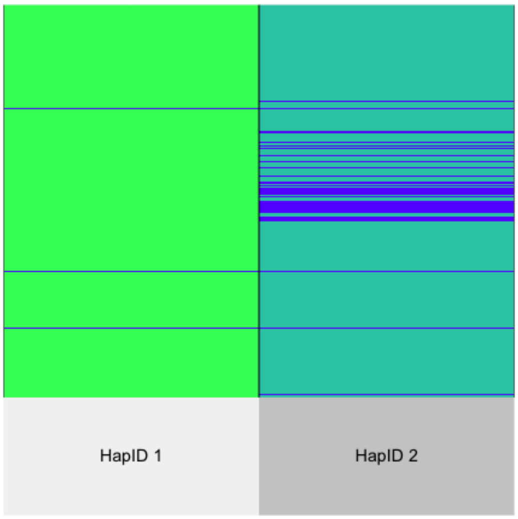


S28a *Structure*

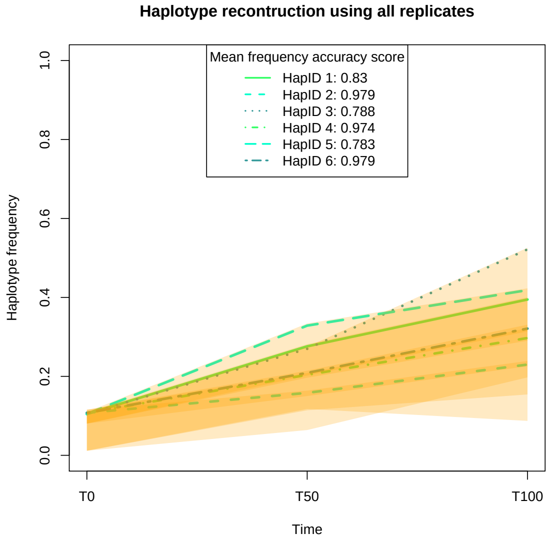


S28b *Frequency*

Figure S28: (a) Match between reconstructed haplotype structure and sequenced founder haplotypes using all the three replicates from [Noble et al., 2019] at the same time. Blue lines indicate mismatches. (b) Reconstructed haplotype frequencies with accuracy intervals (in yellow) and mean accuracy scores using all the three replicates from [Noble et al., 2019] at the same time.

### S12 Results from the HIV data set from [Zanini et al., 2015]

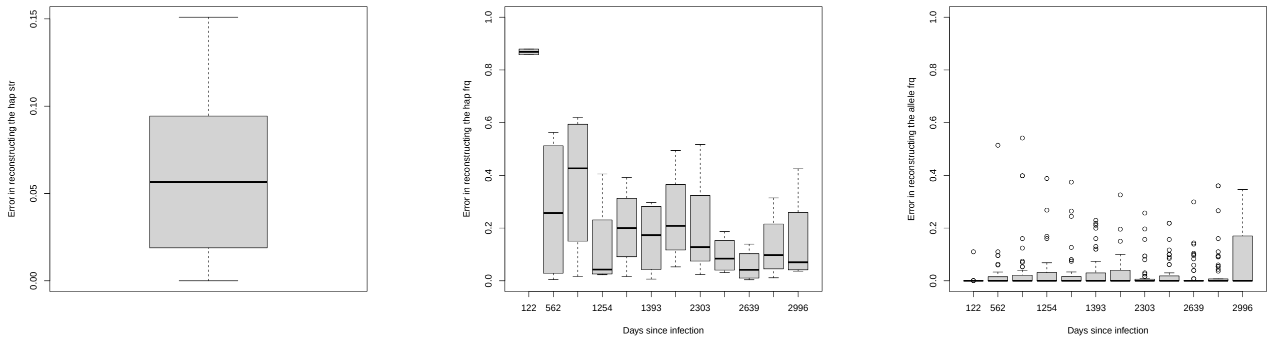


S29a *Structure* S29b *Frequency* S29c *Allelefrequency*

Figure S29: Errors in reconstructing the haplotype structure (left), haplotype frequency (center) and allele frequency (right) for `haploSep` applied to our real data set from patient 1 of [Zanini et al., 2015] and compared on the vpu region.

### S13 Comparison with other methods

In this section we present more details on our method comparison in Section 5. For this purpose we carried out simulations for HIV scenarios. Furthermore, a real data example has also been considered. In this context, we compared `haploSep` with `CliqueSNV` on both the simulated and real data. Unfortunately, we could not obtain results for `CliqueSNV` for all simulated data sets since `CliqueSNV` crashed for 10 samples that were generated in 6 of the 20 simulation runs. We hypothesize that this occurred due to memory problems caused by a large number of candidate haplotypes (30GB memory were allocated to the task). We also considered a comparison with the read based method `PoolHapX` from [Cao et al., 2020]. For our simulated data we could not obtain any results for `PoolHapX`, again because of memory problems with the initial graph coloring part of the algorithm. Nevertheless, we were able to run `PoolHapX` on a very short genomic segment taken from our *D. simulans* E&R real data example. All comparisons have been done using standard model selection provided by the methods to choose the number of reconstructed haplotypes. Further details on the simulation setup are given in Section S13-1 and details on the results are presented in Sections S13-2, S13-3, and S13-4.

#### S13-1 Simulation design

We simulated the data for this comparison using SLiM [Haller and Messer, 2019] using the simulation scenario considered in [Cao et al., 2020]. We followed the script sweep_200loci.slim (`PoolHapX` 1.0.0, downloaded 2/11/2020), but adapted this script to simulate 24 populations. We ran the simulations for a chromosome of length 9719 base pairs, the genome length of HIV. The reads were then aligned to an HIV reference genome available in the `PoolHapX` repository. We simulated 10,000 haploid generations under a sweep scenario. For further information see the SI in [Cao et al., 2020]. From the simulated data we extracted the true haplotype structures and frequencies, as well as the allele frequency data. As input data for `CliqueSNV`, we simulated reads matching the data following the pipeline from [Cao et al., 2020], and we used DWGSIM version 0.1.13 [Homer, 2010] to simulate the reads. `haploSep` has been applied to allele frequency data extracted from these reads.

#### S13-2 Simulations results

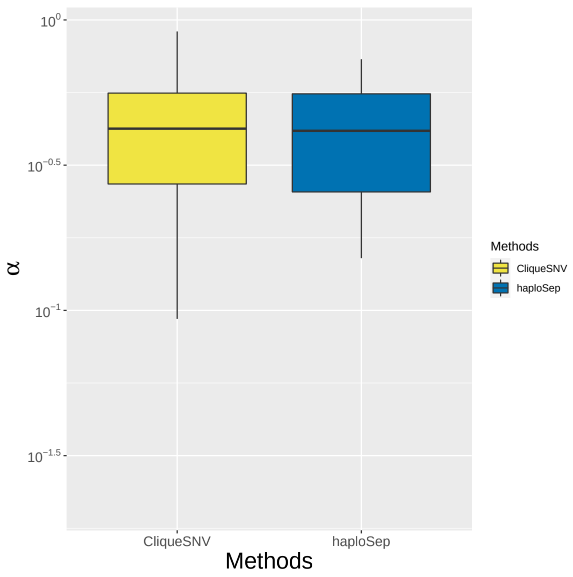


Figure S30: Error ratio ($\alpha$) between haplotype based allele frequency estimates (numerator) and the pool sequencing estimates (denominator) plotted on a log-scale for `haploSep` versus `CliqueSNV`. See Section 3.2 for the definition of $\alpha$. All SNP positions where the observed minor allele frequency is larger than 0.05 in all samples have been used. These results have been derived using the same simulated data as for Fig.5

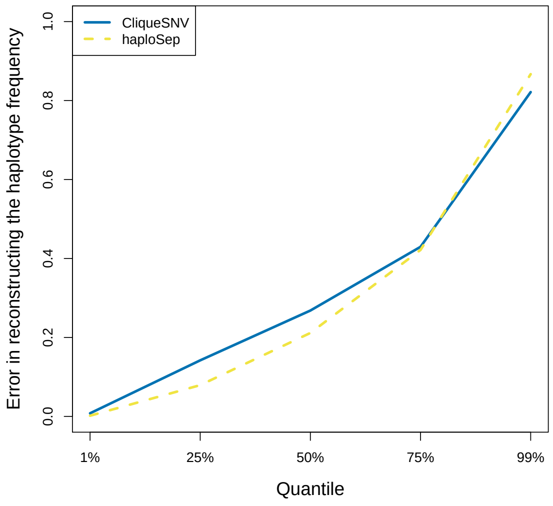


S31a *Frequency*

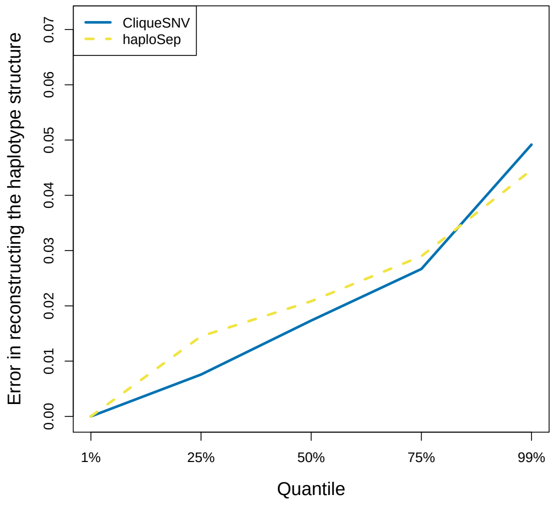


S31b *Structure*

Figure S31: Quantiles of the errors in reconstructing haplotype frequency (a) and structure (b) for both compared methods. The same data were used as in Figure 5.

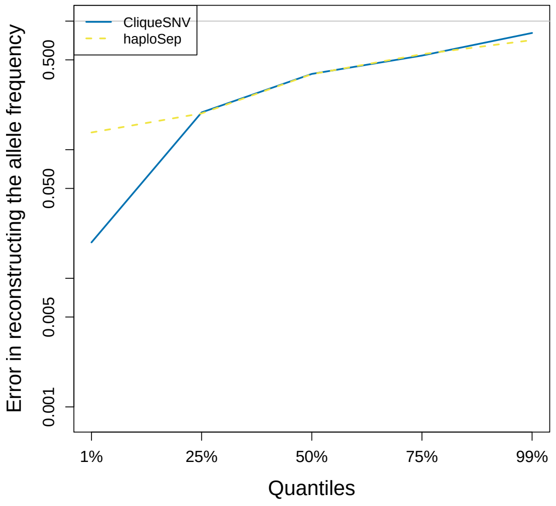


Figure S32: Quantiles of the error ratio ($\alpha$) between haplotype based allele frequency estimates (numerator) and the pool sequencing estimates (denominator) plotted on a log-scale for `haploSep` versus `CliqueSNV`. The same data were used as in Figure S30.

#### S13-3 HIV real data example

Our data consist of 10 time points from HIV patient number 1 in [Zanini et al., 2015]. These data are available in a pre-processed form with the `PoolHapX` package. Our method comparison focuses on the *vpu* window containing 249 SNPs. Besides structure and frequencies, we also compared the sample allele frequencies to the ones reconstructed from the haplotypes and their estimated frequencies using `haploSep` and `CliqueSNV`. For a discussion of Figs. S33 and S34 we refer to Section 5 in the main text. In Fig. S35 we show that `haploSep` leads to fewer outliers, the same median accuracy, and slightly worse 75% percentiles. This is remarkable since this good performance is achieved using fewer haplotypes (4) than the competitor (7–9, depending on the sample), i.e., a less complex model.

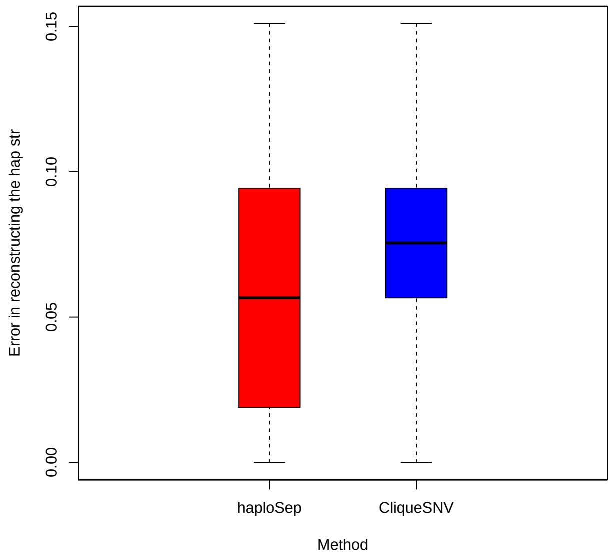


Figure S33: Error in estimating the haplotype structure for `haploSep` (left) and `CliqueSNV` (right) using our real longitudinal data set of patient 1 from [Zanini et al., 2015].

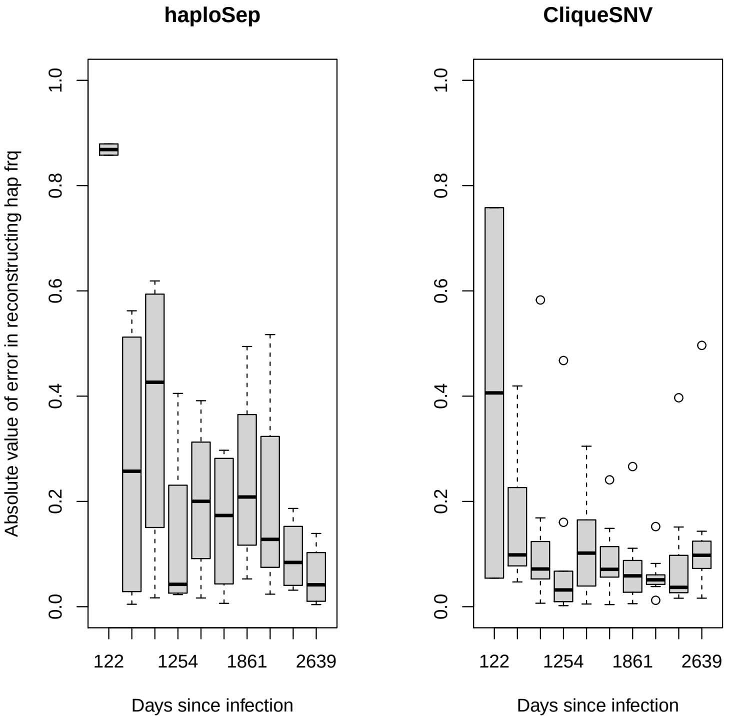


Figure S34: Error in estimating the haplotype frequencies for `haploSep` (left) and `CliqueSNV` (right) using patient 1 longitudinal data from [Zanini et al., 2015].

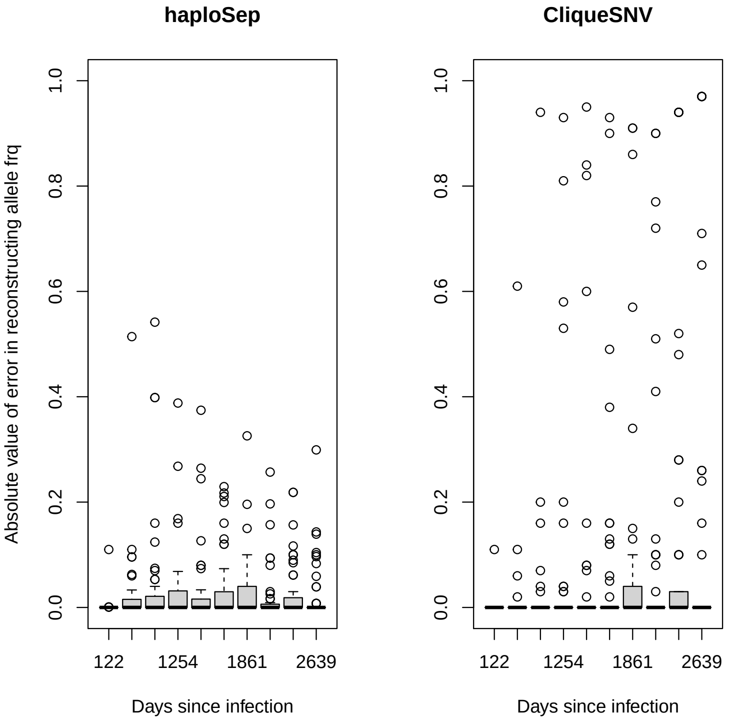


Figure S35: Error in estimating the allele frequencies for each SNP and each time point for `haploSep` (left), `CliqueSNV` (right) using patient 1 longitudinal data from [Zanini et al., 2015].

#### S13-4 *Drosophila simulans* real data example.

We considered a 2200bp region of the *Drosophila simulans* genome containing 95 SNPs, and took our data from [Barghi et al., 2019] (chromosome 2L:11419333-11421533). We tried to apply `CliqueSNV` on this data set using the -os and -oe options to select the above mentioned chromosomal region from the FASTA files provided as reference genome. We obtained run time errors however, indicating that the (20GB) memory of our machine was insufficient. We tried to progressively decrease the window size, but were not able to resolve the problem.

Since `CliqueSNV` did not run for this data, we applied `PoolHapX` instead. `PoolHapX` required 2631 minutes to finish on this small data set. Since `PoolHapX` filters SNPs according to their minor allele frequency (SI of [Cao et al., 2020]), we used the same set of remaining SNPs when we applied our method. As an input to `haploSep` allele frequencies were computed from the available ".bam" files using samtools and Popoolation2 [Kofler et al., 2011].

As the founder haplotype sequences are available in this experiment, we compared the reconstructed haplotypes from both methods with the 189 founder sequences. For each method we consider the most similar founder haplotype and report the percentage of SNPs of the reconstructed haplotypes being identical to the founder sequences. `haploSep` reconstructs three haplotypes and the proportions of matching SNPs are 0.64, 0.48, and 0.72. On the other hand `PoolHapX` reconstructs 21 haplotypes typically less accurately, with proportions of matching SNPs ranging from 0.48 to 0.60.

We also investigated the goodness of fit between the observed and predicted allele frequencies for the competing methods. Fig. S36 shows that the product of the reconstructed haplotype structures with their estimated frequencies provides a closer match to the observed allele frequencies with `haploSep`. It is worth noting that this better fit is achieved based on a much less complex model that uses 3 haplotypes compared to 21 with `PoolHapX`. Therefore the better fit cannot be explained by overfitting.

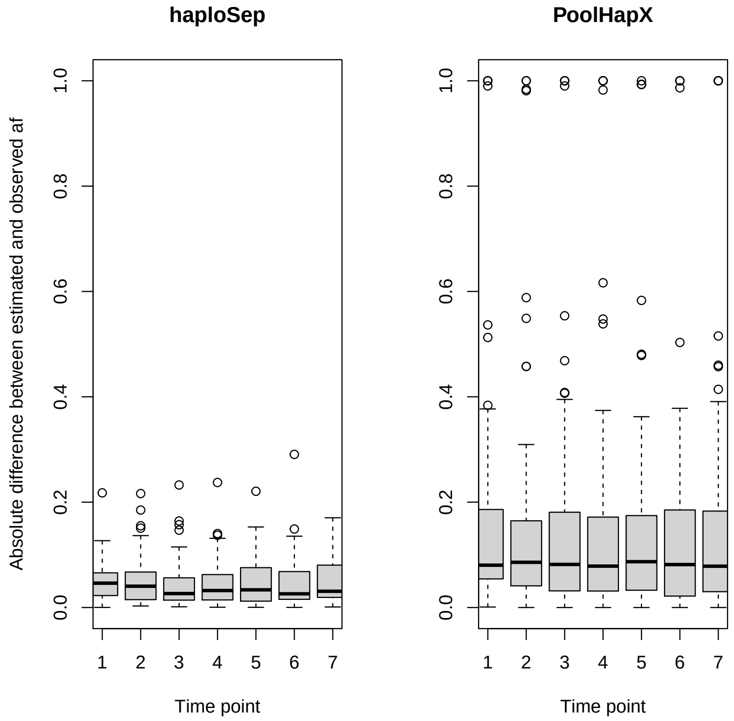


Figure S36: Error in estimating the allele frequencies for each SNP and each time point for `haploSep` (left) and `PoolHapX` (right) using [Barghi et al., 2019] data.

From a computational point of view, there is a striking difference in run time between these two methods. Indeed, the run time for `haploSep` was 0.572 seconds here, whereas for `PoolHapX` the run time was almost 2 days (2631 minutes) on a Mac Pro (2013) machine with 2,7 GHz 12-Core Intel Xeon E5 Processor.
